# Morphological pseudotime ordering and fate mapping reveals diversification of cerebellar inhibitory interneurons

**DOI:** 10.1101/2020.02.29.971366

**Authors:** Wendy Xueyi Wang, Julie L. Lefebvre

## Abstract

Understanding how diverse neurons are assembled into circuits requires a framework for describing cell types and their developmental trajectories. Here, we combined genetic fate mapping and pseudo-temporal profiling to resolve the diversification of cerebellar inhibitory interneurons based on morphology. The molecular layer interneurons (MLIs) derive from a common progenitor but comprise a diverse population of dendritic-, somatic-, and axon initial segment-targeting interneurons. MLIs are classically divided into two types. However, their morphological heterogeneity suggests an alternate model of one continuously varying population. Through clustering and trajectory inference of 811 MLI reconstructions at maturity and during development, we show that MLIs divide into two discrete classes but also present significant within-class heterogeneity. Pseudotime trajectory mapping uncovered the emergence of distinct phenotypes during migration and axonogenesis, well before neurons reach their final positions. Our study illustrates the utility of quantitative single-cell methods to morphology for defining the diversification of neuronal subtypes.

## Introduction

Neuronal diversification is essential for the assembly and function of complex nervous systems (Zeng and Sanes 2017; Cembrowski and Spruston 2019; Geirsdottir et al. 2019). The core feature is the divergence of neuronal populations from shared lineages to subtypes with distinctive features, spatial arrangements, and connectivity patterns (Lefebvre et al. 2015). Subtype diversity arises from the interplay of intrinsic genetic programs and extrinsic cues but capturing the underlying mechanisms remains challenging. By profiling neurons across development with single-cell transcriptomics, one can trace cell fate trajectories based on gene expression signatures (Mayer et al. 2018; Mi et al. 2018; Clark et al. 2019; Tiklová et al. 2019; Trapnell et al. 2014). These studies provide insights into the dynamics and complexity of differentiating neural tissues, including lineage relationships, intermediate progenitors, timelines of maturation and the emergence of subtype identities (Trapnell et al. 2014; Saelens et al. 2019). However, charting neuronal diversification requires additional modalities, such as morphology, to pinpoint emergent structural features and to identify relevant developmental factors.

Although morphology as a modality for cell typing is low throughput, it provides intuitive information about the identity, ontogeny and connectivity of neurons (Gouwens et al. 2019; Scala et al. 2020; Wang et al. 2019; Winnubst et al. 2019). New technologies are advancing the resolution and throughput for mapping cell types based on quantitative morphology (Costa et al. 2016; Winnubst et al. 2019; Economo et al. 2016; Sümbül et al. 2014; Markram et al. 2015; Wang et al. 2019). Correspondingly, large-scale morphological surveys are revealing constituent cell types and providing insights into connectivity patterns and circuit organization (Bae et al. 2018; Helmstaedter et al. 2013; Gouwens et al. 2019; Frechter et al. 2019; Jiang et al. 2015). On the other hand, studies focused on single or related cell types can provide detailed views of local anatomical variation and subtype diversification. This approach requires sparse labeling of genetically or lineage-related cell types, which is facilitated by transgenic drivers and viral vectors (Harris et al. 2014; He et al. 2016). Paired with advancements in statistical and computational methods, neuronal subtypes can be effectively parsed using single neuron anatomy, as illustrated for sensory afferents in the skin (Wu et al. 2012), olfactory bulb neurons (Tavakoli et al. 2018), and pyramidal and GABAergic interneurons in the cortex (Wang et al. 2019; Kanari et al. 2019; Gouwens et al. 2019). Drawing fine divisions between subtypes can be challenging however, as heterogeneity and continuous variation within cell types are commonly observed (Cembrowski and Menon 2018; Gouwens et al. 2019; Harris et al. 2018; Muñoz-Manchado et al. 2018; Scala et al. 2020). Therefore, a major question driving studies of neural diversity is whether all cells sort into discrete subtypes, given sufficient sampling and granularity, or if continuous and local variation is a biological feature essential for neural processing. Large-scale, unbiased morphological analyses have yet to be exploited to study how variation arises during development.

Here, we test a pipeline for mapping the diversification of cerebellar GABAergic interneurons using quantitative morphology. Compared to the complexity of forebrain interneurons (Huang and Paul 2019), the cerebellar molecular layer interneurons (MLIs) derive from a single lineage and form simpler, compact morphologies (Palay and Chan-Palay 1974; Sotelo 2015). Nonetheless, MLIs form an anatomically and functionally diverse population that provides the full complement of dendritic-, somatic- and axon initial segment-targeting inhibition onto principal Purkinje cells. Classically, MLIs are divided into two types—basket cells and stellate cells (Palay and Chan-Palay 1974). Basket cells (BCs) are born earlier, populate the lower third of the molecular layer and form a series of perisomatic basket terminals that envelope the Purkinje cell soma. Some BC terminals further specialize into pinceaux formations that align the Purkinje cell axon initial segment (AIS; Buttermore et al. 2012; Sotelo 2015). By contrast, later-born stellate cells (SCs) settle in the upper molecular layer where their axons innervate Purkinje cell dendrites. However, this classical division has long been debated due to continuous morphological variation observed among MLIs (Sotelo 2015; Rakic 1972; Sultan and Bower 1998). As suggested by fate mapping and heterochronic transplantation studies, the laminar gradient of BC-to-SC phenotypes is related to birthdate and the inside-out settling of precursors within the cerebellar cortex (Altman and Bayer 1978; Cameron et al. 2009; Leto et al. 2009; Sudarov et al. 2011). Terminal commitment into BC or SC phenotypes remains plastic until the final laminar position is reached, suggesting that MLI precursor fates are specified by positional cues (Leto et al. 2009).

In this study, we combined large-scale labeling and reconstruction of MLIs with clustering and pseudotime trajectory inference methods to define the diversity of MLI subtypes at maturity, and to chart the diversification of post-mitotic precursors during development. We devised a novel single MLI anatomy platform that: 1) employs genetic tools to sparsely label MLIs across the repertoire of phenotypes, birthdates and laminar positions through development; 2) reconstructs dendritic and axonal morphologies of MLIs; 3) quantifies phenotypes using morphometric parameters that describe dendritic, axonal, and somatic attributes of each cell; and 4) applies unsupervised clustering and pseudotime ordering algorithms to parse morphological trends among a seemingly heterogeneous neuronal population (Figure 1A). We compiled a dataset of 79 complete reconstructions of mature MLIs, and 732 axonal reconstructions that capture MLIs across development. Our analyses reveal that mature MLIs sort into two discrete subtypes, one of which bears canonical descriptions of basket cells and the other which displays a continuum of basket-stellate cell phenotypes. In a novel approach to quantify diversity based on morphogenesis, we adapted pseudotime algorithms recently developed for single cell RNA sequencing studies to delineate the developmental segregation of MLI identities. In contrast to prevailing views that MLI phenotypes segregate as a function of birth order and laminar position, we provide evidence that basket and stellate cell fates emerge early in their developmental trajectory. Our study also presents a novel framework that is broadly applicable for defining the trajectory of neuronal diversification using morphological information.

**Figure 1:**
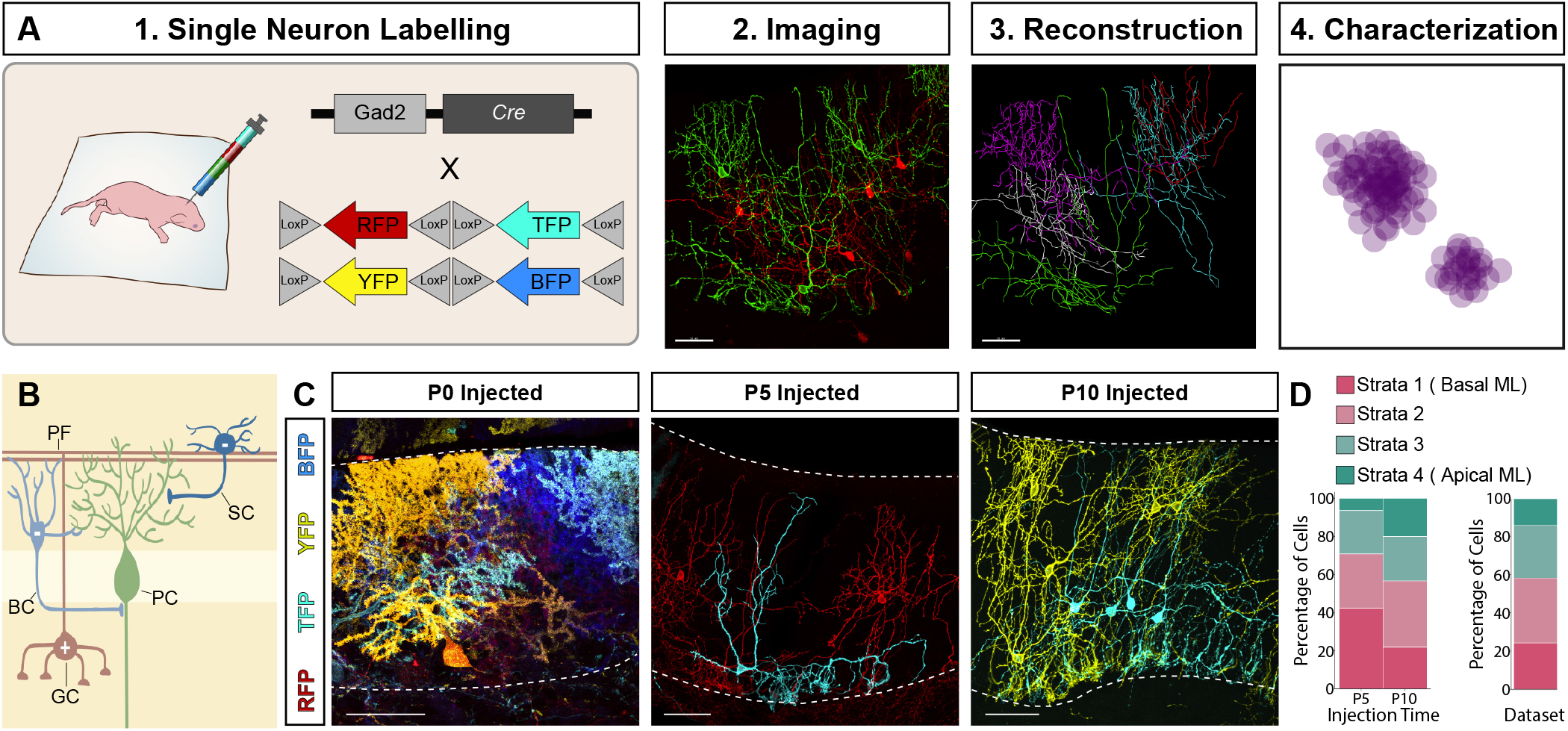
Platformfor morphologicalstudies of molecular layer interneurons. **(A)** Schematic summarizing pipeline for morphological characterizations: 1) single neuron viral labeling with Cre-dependent Brainbow AAVs encoding multiple fluoresce nt proteins (RFP, YFP, BFP, TFP); 2) confocal imaging; 3) semi-automated digital reconstruction and morphometric quantifications using Imaris; 4) multivariate analyses. Scale bars are 50 um. **(B)** Schematic or basket cells (BC, light blue) and stellate cells (SC, dark blue) that reside in the lower and upper moleculau layes (ML), respectively Canonical BC axons target the Purkinje cell (PC) soma and axon initial segment (AISS. SC axons tasget the PC dendrites. **(C)** Inrages of matune cerebellar cortex shows nuorescent labeling of Gad2-Cre positive populations by AAV delivery at different postnatal ages. AAV injection at P0 predominhntly labels PCu; P5, labels lower MLIs; P10, enbiches labelling of upper MLIs. The upper and lower bounds of the ML are outlined by dashed lines. Scale bars ane 50 um. **(D)** Leg: percent laminae distribution ofsingly labeled MLIs in our Brainbow AAV in-ection suheme, based on their laminar distribu)ion defined by soma location in one of four ML strata. Right: Percent laminar distribution of the 79 mature single MLI reconstructions in our dataset.

## Results

### A genetic platform for large-scale labeling of MLIs reveals diverse morphologies

To systematically define MLI diversity, we sought to acquire a high-quality inventory of morphological reconstructions that captures the repertoire of mature MLIs across molecular layer (ML) positions. We devised a single neuron anatomy pipeline in which sparse labeling of MLIs is achieved by injection of *Gad2-ires-Cre* mice with AAV vectors encoding multicolor fluorophores (Cai et al. 2013; Taniguchi et al. 2011). As Gad2-Cre is expressed by other GABAergic populations in the cerebellum, selective MLI labeling was achieved by injecting AAVs at later postnatal stages. Although AAV delivery at P0 resulted in the nearly exclusive labeling of Purkinje cells, injections at P5-7 or P10-14 led to enriched labeling of MLIs residing in the deep or superficial ML, respectively (Figure 1B-D). Images of single MLIs with complete axonal and dendritic arbors were acquired by confocal microscopy and were reconstructed for feature extraction and analyses. We compiled 79 reconstructions of mature MLIs with somas located across laminar positions (Figure 1D), and extracted 27 features that quantify their dendrites, axon, soma, and location (Supplementary Table 1). Together, our dataset contains high-resolution anatomical information of MLIs sampled across ML locations.

We began with manual expert classification of the reconstructions to determine the proportion of MLIs in our dataset that conform to canonical basket cell (BC) or stellate cell (SC) characteristics (Sultan and Bower 1998; Amat et al. 2017). Canonical BCs were identified by soma location in the basal third of the ML, a fan-shaped dendritic arbor that reaches the apical ML, and a long axonal projection which forms multiple basket terminals (16 of 79 cells, Figure 2A, B). Cells with stereotypical SC features were identified by their location in the superficial ML and their highly branched, radial dendritic arbors (25 of 79 cells, Figure 2A, C). Surprisingly, nearly half of the MLIs in our dataset do not fit into either category (non-canonical MLIs, 38 of 79 cells). They include MLIs located in the middle of the ML displaying graded mixtures of BC and SC morphologies that vary with laminar position (Figure 2D), as described previously (Sultan and Bower 1998; Rieubland et al. 2014). Additionally, we uncovered MLIs with morphologies that do not correlate with ML depth. For example, MLIs lacking basket formations but resembling SCs were observed in the deeper third of the ML (Figure 2E), and basket-forming cells were detected in the superficial ML (Figure 2F). Thus, qualitative assessments reveal extensive heterogeneity in axonal and dendritic morphologies among MLIs located across laminar positions.

**Figure 2.**
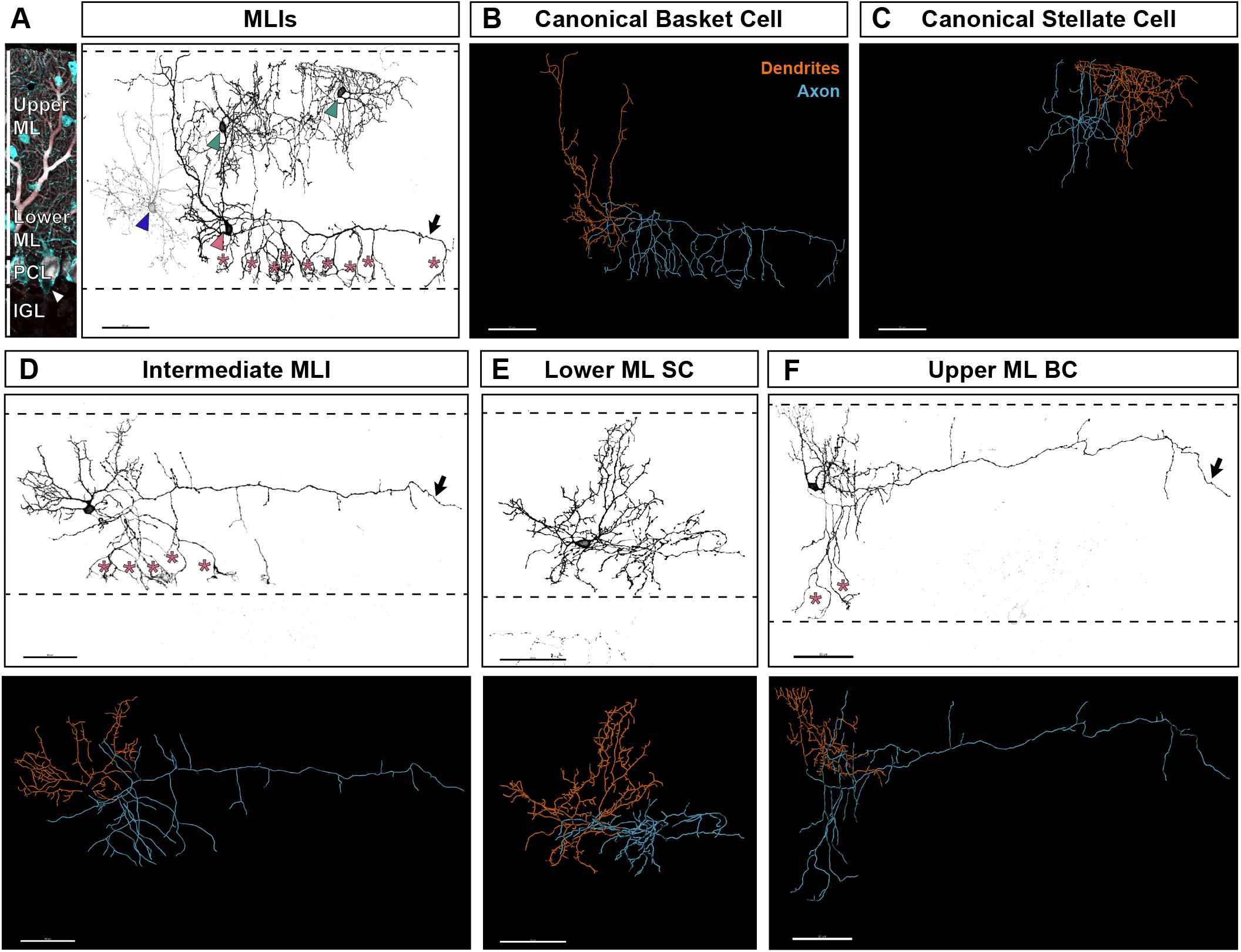
Morphological heterogeneity among MLIs. **(A)** *Left:* Immunostaining of MLIs (Parvalbumin, cyan) and PCs (co-labeled by Parvalbumin and Calbindin, cyan and red, respectively) show the ML space in which MLIs reside. PC somata are enveloped by BC axons (white arrowhead). *Right*: Inverted image of four fluorescently-labelled MLIs. The lower MLI (pink arrowhead) has canonical BC morphologies, including axonal basket terminals around the PC somata (asterisks). The two upper MLIs (teal arrowhead) have canonical SC morphologies and reside within the upper ML. The faintly labelled MLI (blue arrowhead) has SC morphologies but reside within the lower ML. **(B)** Reconstruction of canonical BC from panel A, with complete dendritic arbor traced in ofange, and complete axonal arbor traced in blue. **(C)** Reconstruction of canonical SC from panel A (cell on far right). **(D-F)** Redresentative images of MLIs with mixtures of BC and SC characteristics. *Top:* inverted fluorescence image; *Bottomr* reconstruction with dendritic (orange) and axonal (blue) traces. **(D)** MLI located in the middle ML with SC dendritic features, end a long horizontal axon (arrow) with descending basket collaterals (asterisks,). **(E)** MLI located in the lowe (ML with SC-like dendritic and axonal arbors. **(F)** MLI located in the upper ML with a long horizontal axon (arrow) and two descending axon collaterals, with basket Oormations around PC nomas (asterisks). All scale bars are 50 μm.

### MLIs form two discrete cell types that are distinguished by axonal signatures

To test if the repertoire of MLIs can be distinguished into subtypes, we applied unsupervised clustering methcds to analyze the dataset of single MLI morphometric properties. We first performed hierarchical clustering which rendered two major clades figure 3A). We inspected each reconstruction to relate the hierarchical ordering to morphological phenotypes. The first clade contained basket cells displaying canonical features and corresponding to the manually curated BCs (n = 19 cells; Figure 3A-C). By contrast, the second clade was larger, comprising canonical SCs and the remaining non-canonical MLIs (n = 60 cells; Figure 3A). Notably, cells in this SC-like clade (hereafter SCs) were ordered into subclades that followed a linear progression in morphological characteristics. Visually, we discerned four subdivisions that reflected similarities in shapes and ML locations: SC1, stellate cells with short-range axons located in the bottom ML (Figure 3B, D; non-basket forming); SC2, short-range stellate cells in the middle and upper ML (Figure 3B, E); SC3, stellate cells in the middle and upper ML with long-range axons, some of which give rise to 1-3 basket collaterals (Figure 3B, F); and SC4, long-range stellate cells constrained to the apical ML (Figure 3B, G). By statistical methods however, the optimal number of clusters for the total MLI dataset were identified as two by silhouette analysis and to three by the elbow method (Supplementary Figure S1; Rousseeuw 1987; Thorndike 1953). We reasoned that further divisions may be ambiguous due to the limitations of these tests or due to heterogeneity among the SC population.

**Figure 3.**
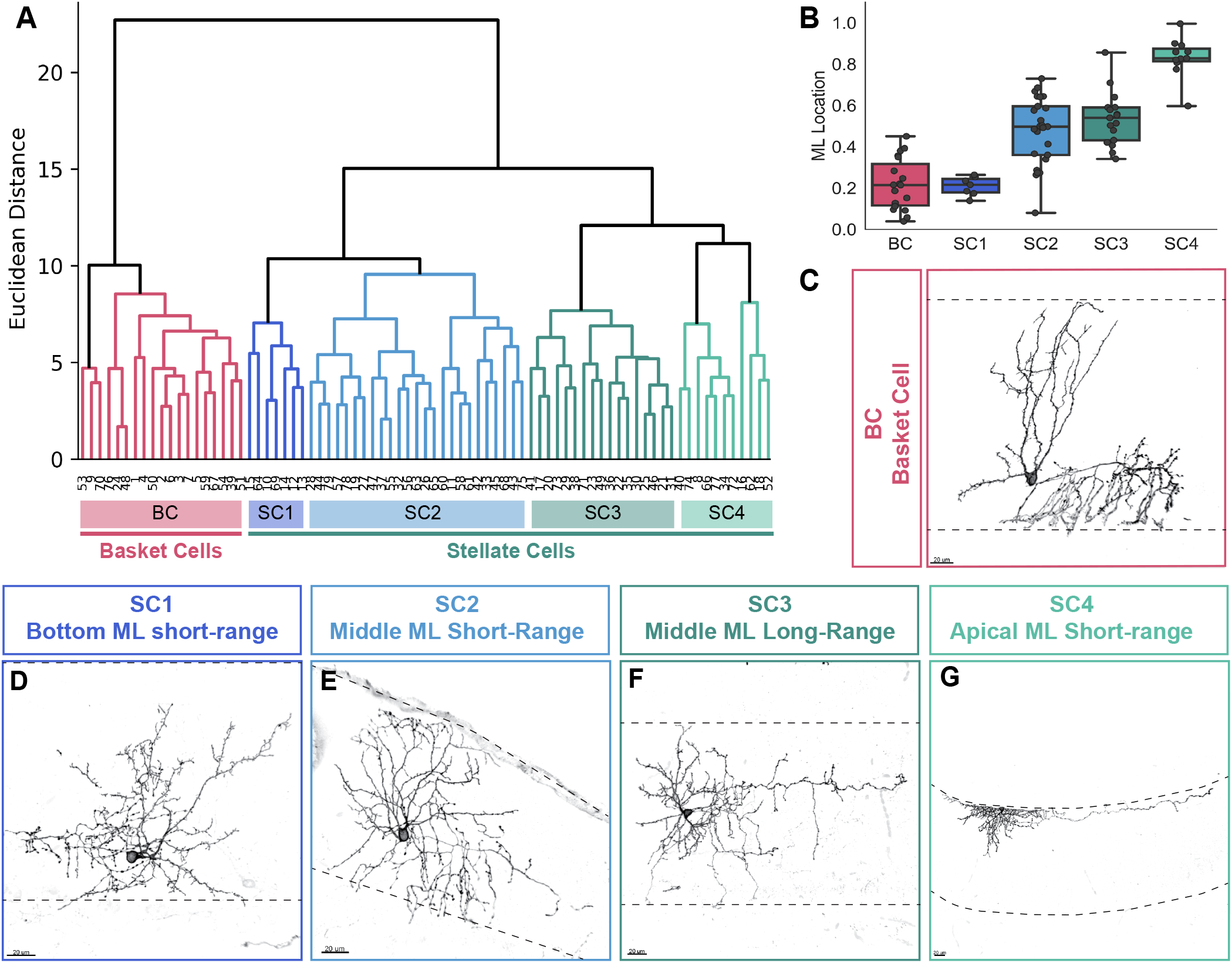
Hierarchical clustering of mature MLI morphologies unmasks two distinct morphological groups. **(A)** Hierarchical clustering dendrogram. The BC cluster (pink) is separate from the SC cluster (blue-greens). Within the broad SC cluster, the dendrogram ordered MLIs based on shared morphological characteristics and was mgnually curated into foursubclades based on expert assessment or similarities. **(B)** Ml_ Location of MLIs sorted into designated subclades. **(C-G)** Individual MLI examples from the SC subclades (SC1-SC4). Dashed lines denote the apical and basal extremities of the Ml_. Scale bars are 20 μm.

To test if MLIs could be further parsed into subtypes by other clustering methods, we applied the Uniform Manifold Approximation and Projection algorithm to the morphometric dataset (UMAP; Becht et al. 2018; McInnes et al. 2018). UMAP is a dimensionality reduction algorithm that projects each MLI reconstruction onto a two-dimensional space based on local similarities with other MLIs. UMAP rendered two clusters and reproduced the BC and SC division, similar to hierarchical clustering (BC cluster, n = 18 cells; SC cluster, n = 61 cells; Figure 4A-F). Although the SC cluster is broad and contains morphologically heterogeneous cells, the cells largely co-distributed within the curated phenotypic subclades from hierarchical clustering (Figure 4B, E, F). Moreover, distantly projected cells exhibited different phenotypes, such as the projection of lower ML short-range SCs at one end of the cluster and upper ML long-range SCs at the other (Figure 4 D,E). A similar division between the BC and SCs was produced with the t-Distributed Stochastic Neighbor Embedding (t-SNE) algorithm (Supplementary Figure S2). Taken together, these analyses robustly divide MLIs into two discrete populations, the BC and SC types, but the SC population exhibits considerable heterogeneity.

**Figure 4.**
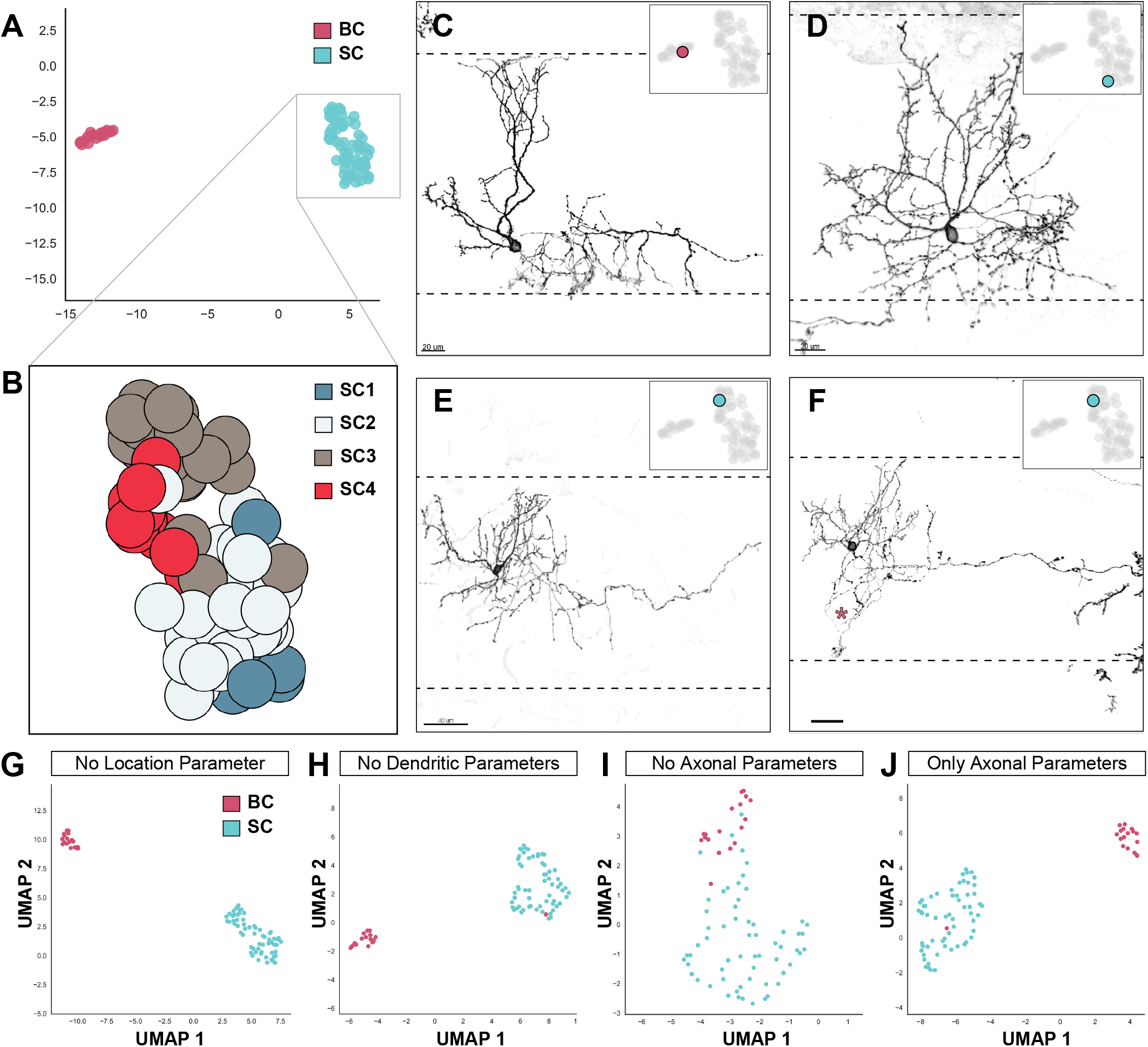
UMAP of mature MLI morphologies uncovers two distinct subtypes. **(A)** UMAP projection of 79 mature MLI reconstructions in our dataset. Colors indicate BCs (pink) and SCs(teal). **(B)** Projection of SC subclade identity from hierarchical clustering tesults projected onto the SC cluster. **(C-F)** Individual examples of MLIs, validate the morphological correspondence between MLI clusters. **(C)** MLIs from the BC cluster (inset) display canonical BC moyphologies. Basal and apical limits of the ML are outlined by dashed lines. **(D)** An MLI at one pole of the SC cluster (inset) displays short-range axonal morphology and resides within tire basal ML. **(E)** An MLI at the opposite end (inset) displays long-range axonal morphology, while a cell at a neighboring datapoint **(F)** has a similar Corm but extends descending axon collaterals to PC somata (pink asterisk). **(G-J)** UMAP projections following recursive elimination of morphological features describing ML location **(G)**, dendrites rtt), or axons **(I)**. Axonal information was necessary **(I)** and sufficient (J) foy UMAP-dependent MLI clustering. Scale bars are 20 μm for C, D and F, and 40 μm for E.

We next examined the morphological within-class heterogeneity present within the SC population. It was previously proposed that MLIs model a single continuously varying population based on ML laminar positioning (Rieubland et al. 2014; Sultan and Bower 1998). We wondered if this heterogeneity may be accounted for through variations within the SC cluster alone. To test this possibility, we performed regression analysis on all morphological features as a function of ML location, first using all 79 reconstructions in our dataset (Supplementary Figure S3). We found 8 parameters which showed significant positive or negative correlation with soma position within the ML, including dendritic complexity, as previously described (Rieubland et al. 2014). We performed an additional round of regression analysis for each of these 8 parameters, using only the 61 SCs in our dataset. We found that for 4 of these parameters, the correlation remained even after removal of all BCs from the analysis (Supplementary Figure S3). The remaining 4 parameters corresponded to features where the BC population added a significant skew to the correlations graph, and where the entire population had not modeled continuous variation in the first place. We further confirmed that all 4 parameters followed a largely normal distribution within the SC type, with no presence of bimodality (Supplementary Figure S4). Together, our results suggest that the significant variation among SCs denotes continuous variation, similar to what was proposed previously for the entire MLI population.

Finally, to define the features most informative for MLI classification, we analyzed the dataset using recursive feature elimination, in which single or groups of morphological features were removed. The BC/SC division remained following elimination of all dendritic features and, surprisingly, elimination of soma location (Figure 4G, H; Supplementary Figure S2). By contrast, BC and SC clustering was lost upon removal of all axonal features (Figure 4I). To test whether axonal phenotypes alone are sufficient for clustering, we restricted UMAP analyses to six parameters that describe axonal morphology. Interestingly, the BC and SC clusters were recapitulated with all but one cell correctly sorted (Figure 4J). Therefore, despite the anatomical complexity and heterogeneity, axonal information is necessary (Figure 4I) and sufficient (Figure 4J) for the classification of MLIs into BC or SC subtypes. We conclude that MLIs form two discrete cell types. While our results support the classical BC and SC division, the significant heterogeneity present within the SC group also explains previous observations of continuous variation that was the basis for the one MLI population model (Sultan and Bower 1998).

### Genetic fate mapping of MLIs reveals a population of early-born stellate cells

Having established the divergence of MLIs into two subtypes on the basis of axonal information, we wondered whether the emergence of MLI identities could be traced during development based on axon morphologies. The differentiation of MLIs is largely uncharacterized and there are no established molecular markers to distinguish BCs and SCs at maturity nor during development (Sotelo 2015; Schilling and Oberdick 2009). Therefore, we reasoned that morphological mapping could reveal time points and locations that influence MLI diversification into basket and stellate cell phenotypes. To date, fate mapping and heterochronic cell transplantation studies have led to a series of observations that together suggest that MLI identities are determined by birth order and laminar location. First, MLIs are generated from a shared progenitor pool but laminar specific phenotypes result from the sequential generation and the inside-out settling of precursors within the cerebellar cortex (Altman and Bayer 1997; Leto et al. 2009; Sudarov et al. 2011; Zhang and Goldman 1996b). Second, terminal commitment into BC or SC identities is plastic until the final laminar position is reached (Leto et al. 2009). According to this model, early-born MLIs that first populate the lower ML adopt BC identities, while later-born MLIs that fill the upper ML become SCs. To directly test this model, we devised a genetic fate mapping strategy to label single MLIs born on different days and trace their locations and patterns of axon morphogenesis. We used the tamoxifen-inducible *Ascl1-CreER* line to activate Cre in MLI progenitors at the time of terminal division (Sudarov et al. 2011; Brown et al. 2019). Consistent with previous studies, tamoxifen (TMX) injection of Ascl1-CreER; Ai4^flox-STOP-TdTomato^ animals at P0 enriched for the labelling of BCs that innervate the lower ML, while delivery at P4 and P7 marked SC-like cells restricted to the upper ML (Figure 5A-C).

**Figure 5.**
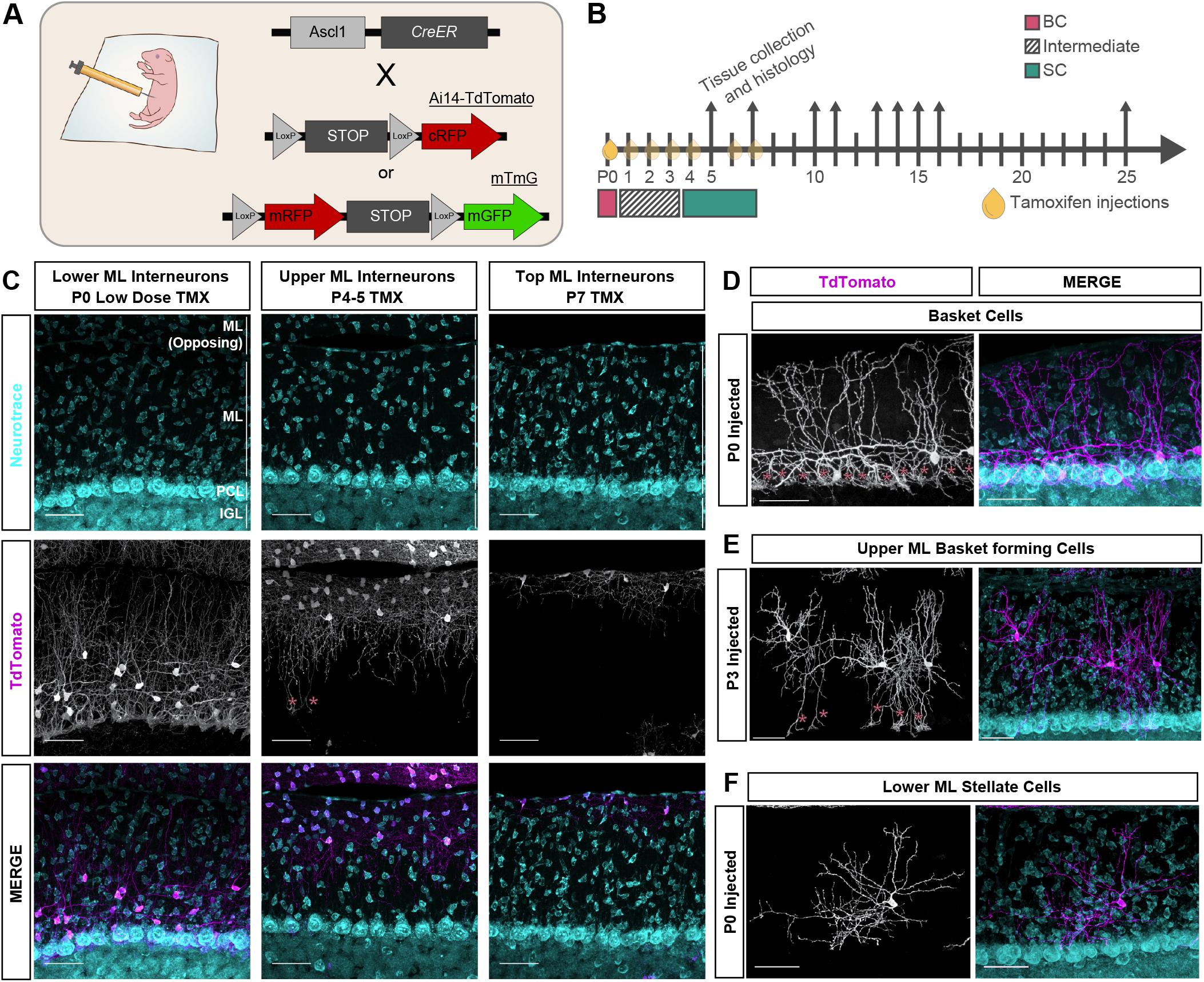
Birthdate-dependent targeting of MLI subpopulations by Ascl1-CreER. **(A)** MLI labeling strategy by tamoxifen (TMX) inducible Ascl1-CreER and fluorescent Cre reporters, Ai14-TdTomato (cytosolic) or mTmG (membrane-targeted). **(B)** Schematic for TMX-induced subtype-enriched MLI labelling. Postnatal timepoints of single dose TMX injections are depicted by oil droplets. Collection timepoints are shown by upwards arrows. Lowdose induction at P0 predominantly labels BC subpopulations, P2-P3 induction labels intermediate MdIs, P4-7 indections label SC cells in the upper ML. **(C)** Confocal images of Ascl1-CreER; Ai14 cerebellar cortex at P25 shows laminar restriction ofMLI labelinc) (TdTomato, greyscale or magenta; Neurotrace, cyan) following TMX induction ad P0, P4, or P7. **(D)** Example of P0-induced BCs in the lower MC. **(E)** Example of P3-indfced SCs in the upper ML with descending baeket terminals. Pink asterisks denote basket formations enveloping PC soma in D, E. **(F)** Example of P0-induced SC in the lower ML. Representative images were taken at P25. Scale bars are 50μm.

To determine the relationship between MLI birthdate and terminal fate, we characterized the mature morphologies and laminar positions of MLIs labelled by TMX injections on different days. Additionally, we aimed to relate SC subclade identities established from hierarchical clustering analyses to their birth order (Figure 3). CreER induction at P0 labelled BCs in the deep ML (Figure 5D), while induction at P2-3 predominantly marked MLIs with mixed BC-SC phenotypes, consistent with the middle and upper ML basket forming cells of SC3 (Figure 5E). The majority of MLIs labelled at P4-5 were non-basket forming short- and long-range SCs in the superficial ML, typical of SC2 and SC3 (Figure 5C). P7 inductions labelled apical ML constrained SC4 cells nearly exclusively (Figure 5C). Intriguingly, we observed that SC1s which reside within the lower ML were marked at P0, and to a smaller extent at P1 (Figure 5F). However, they were not detected following CreER induction at P3 and onwards, suggesting that the lower ML SC1s are early-born cells within the MLI lineage. These findings support one aspect of the current model in that the inside-out stratification of MLIs within the molecular layer is temporally related to birth order. However, they also suggest the existence of an early-born SC subpopulation which shares a similar temporal origin and laminar fate to BCs but adopts a distinct, non-basket forming phenotype. Our findings therefore indicate that BC and SC phenotypes are not strictly separated by temporal origin or laminar positioning.

### MLI axon morphogenesis can be ordered along a pseudo-temporal maturity gradient

The marking of early-born MLIs that elaborate distinct phenotypes in the lower ML suggests the divergence of BC and SC fates during early postnatal development. We hypothesized that if MLIs diverge into two early-born BC and SC precursors, the subpopulations would progress through distinct developmental trajectories. Alternatively, if MLI subtype differentiation is dependent on positional cues, such as those present at the site of integration, then MLI precursors might share a similar progression until they settle and elaborate subtype-specific axonal arbors. To distinguish between these possibilities, we sought to define the divergence of BC and SC identities by comparing patterns of migration and axon morphogenesis. To incorporate developmental data spanning all stages of development, we adapted a pseudo-temporal ordering approach to align snapshots of single neuron morphologies along a continuous progression of maturation. Pseudotime trajectory inference was recently developed for analyses of single cell transcriptomics datasets, where cells dissociated from developing tissue at a fixed time contain a spectrum of cellular states due to the asynchronous nature of development (Figure 6A; Setty et al. 2019; Trapnell et al. 2014; Saelens et al. 2019; Serra et al. 2019). Thus, computational alignment of single cell data along a continuous maturity gradient enables a model for the progression and bifurcations of cellular states (Trapnell et al. 2014). In a similar vein, we applied pseudotime modelling to infer the continuous trajectories of MLI development using high dimensional morphometric data of single neurons (Figure 6A).

**Figure 6.**
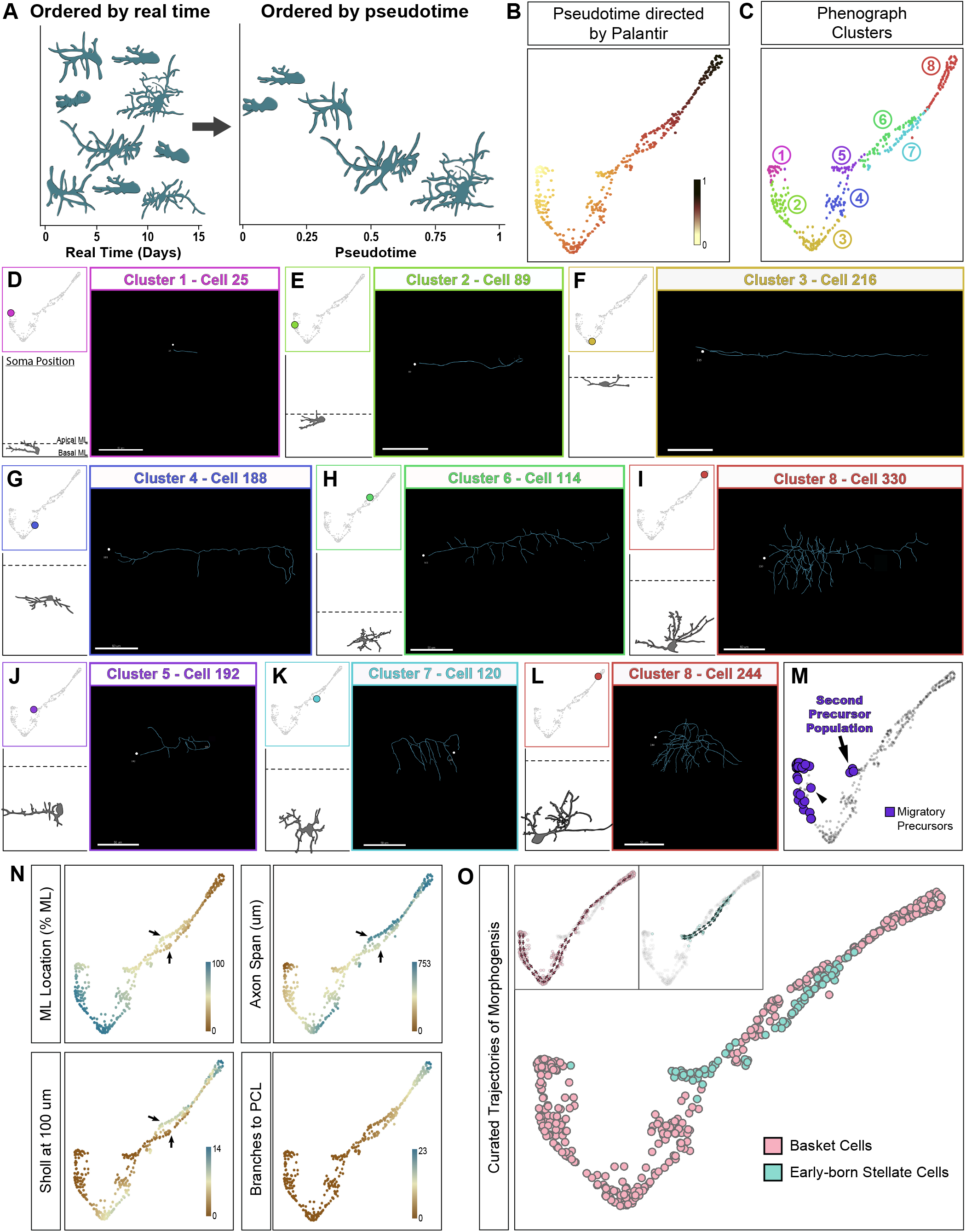
Pseudotime ordering of early-born MLIs based on quantitative analyses of axonal development reveals two lineage trajectories. **(A)** Schematic showing the basis of pseudotime trajectory inference methods. If ordered by real time (i.e. by animal age, *left),* progressions of neuronal maturation are difficult to ascertain due to variability of developmental stages. Pseudotime orders snapshots of single developing neurons based on morphological progression rather than animal age (*right*). **(B)** Palantir-generated pseudotime ordering of early-born MLIs (P0 TMX injection - collected P5 - P25). Each datapoint represents one reconstruction from the dataset of developing MLI axon morphologies. **(C)** Phenograph-generated division of the axonal dataset into eight clusters that reflect different morphological stages. **(D-M)** Examples of single MLI morphologies plotted within each cluster 1-8. *Top left:* Position of cell in pseudotime trajectory. *Bottom left:* Camera Lucida illustration of cell soma and dendrites, positioned to scale by ML location, to validate progressive maturation of cells. *Right:* Axonal reconstruction with soma outlined by white dot. **(D-I)** Representative examples along the BC trajectory of morphogenesis. **(J-L)** represent examples of MLIs along the early-born SC trajectory of morphogenesis. **(M)** Migratory precursors are plotted in two locations. In addition to the precursors for the BC population (cluster 1, black arrowhead), there is a second naive population that corresponds to early-born SCs (arrow). **(N)** Heatmap representation of single morphometric parameters corresponding to Palantir-ordered MLIs. Arrows highlight differences between the BC and early-born SC trajectories. BCs occupy higher positions within the ML during migration (ML location) with axons that span a greater distance along the ML (axon span and Sholl at 100um). **(O)** Curated pseudotime trajectories for early born MLIs. BCs are highlighted in pink, while early born SCs are highlighted in cyan. All scale bars are 50 μm.

We acquired a large dataset of developing MLI morphologies marked by low-dosage TMX injections and collected over a series of timepoints spanning MLI morphogenesis (P5 - P21) (Figure 5B). Axonal arbors of 732 MLIs were reconstructed and quantified for 28 morphometric parameters describing their axon, soma and location. Each reconstruction was annotated for the cell’s inferred terminal fate (early-born, BC or SC1s: P0 TMX injected; late-born, SCs: P3-7 TMX injected) and age (days post TMX injection, hereafter DPI). In parallel, each cell was manually binned into one of four maturation stages according to a qualitative assessment of dendritic arborization and soma shape (Supplementary Figure S5). This scheme served as an expert-directed ordering to validate the performance of pseudotime algorithms in aligning axonal reconstructions along a maturation trajectory.

We first evaluated the suitability of pseudotime algorithms for modeling axonogenesis and if so, to determine whether separate BC- and SC-fated trajectories can be distinguished. We began with analysis of the early-born MLIs (P0 TMX, 423 cells) using Palantir, a diffusion map-based pseudotime algorithm (Setty et al. 2019; Coifman et al. 2005). Palantir rendered a trajectory that followed a linear progression, as confirmed through both the default and expert-directed pseudo-timelines (Figure 6B; Supplementary Figure S6, 7). To evaluate the accuracy of the manifold, we clustered the trajectory into eight developmental states using PhenoGraph (Levine et al. 2015), which computationally partitions high-dimensional datasets into phenotypically meaningful subpopulations (Figure 6C). Inspection of clusters along the trajectory revealed a robustly ordered pseudotime ordering of canonical BC axon morphogenesis, in clusters 1, 2, 3, 4, 6, and 8 (Figure 6C-I). Cluster 1 marked the beginning of the BC trajectory, where cells displayed the simplest morphologies consisting of a single, short axonal extension (Figure 6D). The end of the trajectory was represented by cluster 8, which contained cells with the most complex axonal arbors, and increasing elaborations of basket terminals (Figure 6I). Importantly, the intervening clusters uncovered previously unknown trends during BC morphogenesis, in addition to known characteristics. Cluster 1 and 2 cells possessed a leading process and exhibited immature soma morphologies oriented along the tangential plane of the apical ML (Figure 6D, E), consistent with descriptions of tangentially migrating MLI precursors (Cameron et al. 2009; Wefers et al. 2018; Wefers et al. 2017). This finding suggests that BC axonogenesis begins during neuronal migration. Consistently, cells in clusters 3, 4, 6 and 8 occupied gradually decreasing laminar positions as axonal morphogenesis proceeded to completion (Figure 6F-I). Together, the robust ordering of BC axonogenesis demonstrates the suitability of pseudotime algorithms for aligning snapshots of single neuron morphometric information and inferring the trajectory of morphogenesis.

Notably however, MLIs represented in clusters 5 and 7 do not fit the axonal progression of BCs (Figure 6C). Compared to cells in clusters 3 and 6 with similar extent of dendritic maturation, axonal arbors of cluster 5 and 7 cells have noticeably shorter primary axons (Figure 6J-L). Additional visual curations identified a small population of migratory precursors lacking a noticeable trailing process, a characteristic of immature MLIs that were also mapped to clusters 1 and 2 (Figure 6M). This observation suggests that cluster 5 marks the beginning of a second early-born MLI trajectory (Figure 6M). We further examined this divergence by projecting individual morphometric parameters onto the manifold. The parallel tracks formed by clusters 6 and 7 differed in axonal span, branching complexity and laminar location (Figure 6N; Supplementary Figure S8). A similar but more substantial contrast was noted between clusters 3 and 5 (Figure 6N; Supplementary Figure S8). Taken together with our previous fate mapping studies and given the short-range nature of cluster 5 and 7 cells, our results support a model of two lineages of early-born MLIs that correspond to BCs and the lower ML, short-range SC1s (Figure 6O)

Given the power of pseudotime inference for modeling of early-born MLI morphogenesis and uncovering *de novo* trajectories, we next applied Palantir to the entire suite of 732 MLI reconstructions. Palantir produced a pseudo-timeline but the trajectory was discontiguous, with alternations between immature and mature pseudotime states (Figure 7A). To test if this was due to the altered alignment of cells within the manifold, or incorrect projection of pseudotime states to the underlying manifold, we projected the manually annotated single cell maturation stages onto the trajectory (Figure 7A and Supplementary Figure S5). Our expert-directed pseudotime approach confirmed that the underlying manifold structured the cells from immature to mature states (Figure 7A). We further mapped the BC and SC1 lineages, as well as the late-born SCs (TMX at P4-P7; Supplementary Figure S9). However, the projections were difficult to interpret as cell lineages were arranged along a discontinuous pseudo-timeline and the global lin eagre structures we re not maintained. In an alternate approach, we clustered the trajectory into ten developmental states and analyzed the local subtype distributions (Figure 7B; C). Nine of the ten clusters were enriched for either the early-born or late-born MLI populations, regardless of pseudotime maturity, which suggests clustering at the local scale (Figure 7C). The remaining cluster contained SC1 cells nearly exclusively (Supplementary Figure S10). Although the Palantir-directed trajectory separated MLI subtypes within pseudotime stages, the overall performance with this dataset was limited.

**Figure 7.**
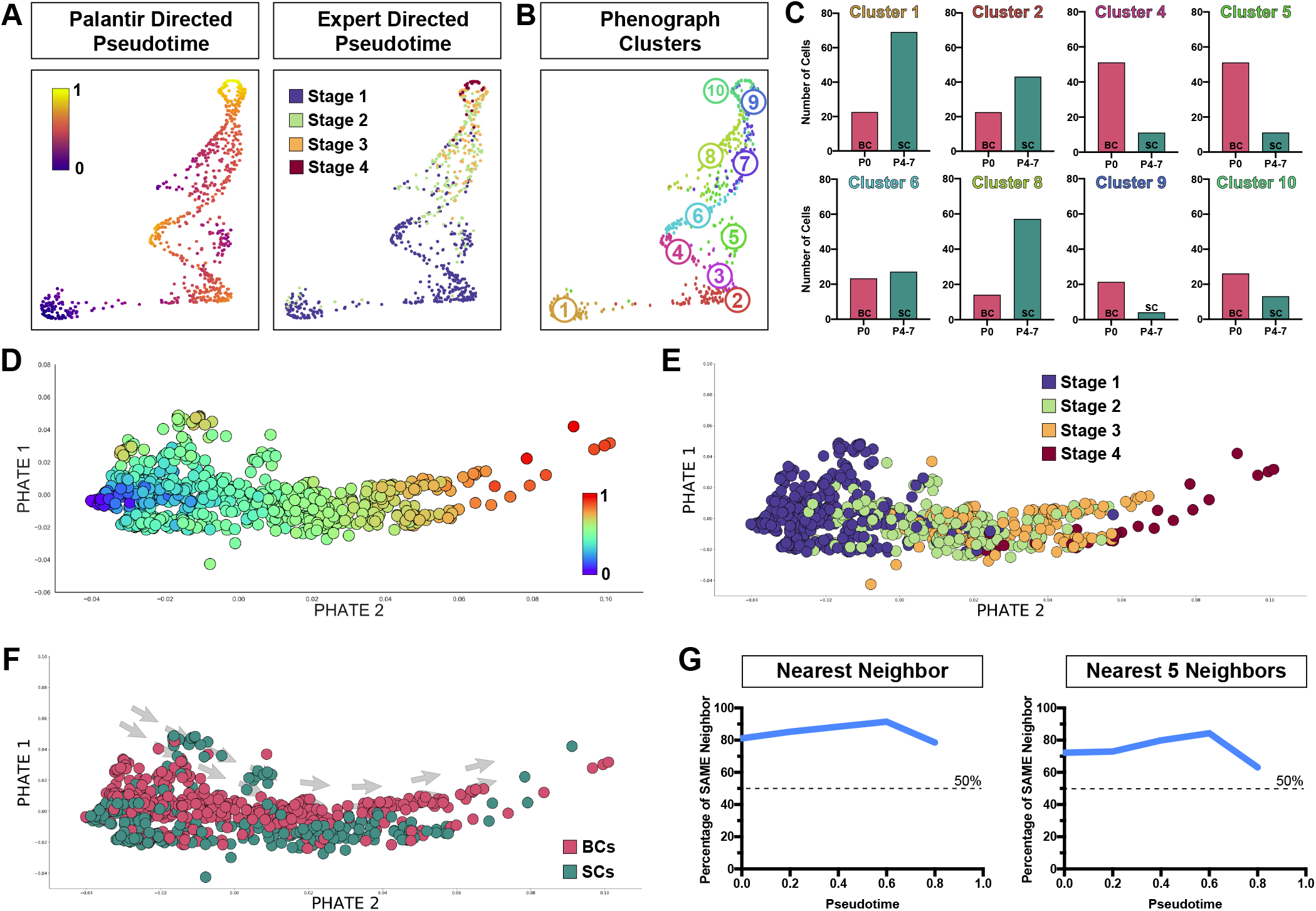
Emergence of MLI-type identities prior to axonogenesis. **(A)** *Left:* Palantir-generated pseudotime ordering of axonal traces from 732 developing MLIs. *Right:* Expert-directed pseudotime ordering of reconstructions using manual annotation of maturation based on 4 stages of dendrite progression. **(B)** Phenograph-generated clustering that reflects different morphological stages. **(C)** Number of BCs (P0 TMX injected) or SCs (P4-7 TMX injected) within each Phenograph cluster. Nearly all clusters show enrichmentof BCs ocSCs. **(D)** PHATE-cenerated pseudotime ordering. **(E)** Validation or PHATE ordering by projecting manual annotation ofdendrity maturity foreach MLI. **(F)** PHATE generated pseudotime trajectory for the entire MLI data set, colored by BC (early born, P0 TMX; pink;) or SC identity (late born, P4-P7 TMX; teal). **(G)** For each neuron (datapoint), the nearest 5 neighbors were obtained. *Left:* proportion of cells with a nearest neighbor of identical fate (i.e. BC vs SC). *Right:* Proportion or nearest 5 neighbors with the same identity.If MLI fates are established late during development, ~50% or cells would be expected to have a neighbor of the same identity (dashed line). Over 80% or MLIs at pseudotime 0 have an identical neighbor, suggesting early establishment of MLI identity. Th e drop in pe rcentage of ide ntical neighbors between 0.6-0.8 pseudotime is l i kely due to sparsity of mature reconstructions in our dataset, i.e. fewer delta points at the right ofthe PHATE trajectory in **(D-F)**.

### Trajectory inference by PHATE reveals the early emergence of BC and SC identities

To test if the total MLI trajectory can be resolved by other trajectory inference algorithms, we applied a parallel approach using PHATE. PHATE was developed for visualization of branching data structures, with preservation of both local and global similarities (Moon et al. 2019). The PHATE-generated trajectory ordered the MLI reconstructions along a linear arrangement that reflected a pseudo-temporal gradient (Figure 7D), and was confirmed by our expert-directed staging metric (Figure 7E). Projection of the early- and late-born MLIs revealed that the two populations largely segregated along the trajectory from the beginning of pseudotime (Figure 7F). To quantify this observation, we performed nearest neighbor analyses following pseudotime binning (Figure 7G). We reasoned that if the terminal fates of MLI precursors remain undetermined at the beginning of axonogenesis, the identity of a cell’s nearest neighbor should follow a largely random distribution. Correspondingly, roughly 50% of early pseudotime cells should possess a nearest neighbor of the same prospective terminal fate. Alternatively, a positive or negative deviation from 50% infers an early bias of MLI type identities. In support of the latter, 80% of MLIs at the origin of pseudotime possessed a nearest neighbor with an identical terminal fate (Figure 7G). We further confirmed this finding by extending the analysis to the nearest 5 neighbors (Figure 7G). Taken together, these results provide evidence for an early divergence model for MLI type identities. Our data support a scenario where migratory MLI precursors at the beginning of axonogenesis are biased towards their terminal fates.

To determine whether divergent axonogenesis patterns are visually discernible, we compared pseudotime to real time morphologies within the *in situ* tissue space. At 5-6 days post injection (DPI), we observed tangentially migrating BCs and SCs that bear a long axonal process (Figure 8A, B). Their morphologies resemble cluster 1 cells of the BC pseudotime lineage and are representative of the stage 1 manual classifications (Supplementary Figure S5). By 10-11 DPI, MLI precursors displayed a mixture of developmental stages that were stratified according to the inside-out layering of the ML (Figure 8C, D). Tangentially migrating stage 1 BCs and SCs were located in the apical ML, while stage 2 and 3 cells were located in progressively lower ML positions corresponding to increasing axonal complexity (Figure 8C, D). The developmental gradient of cells across the ML supports our pseudotime findings (see Figure 6N). By ~20 DPI, nearly all labelled neurons displayed axonal morphologies characteristic of mature BCs or SCs, consistent with the end of pseudotime (Figure 8E, F).

**Figure 8.**
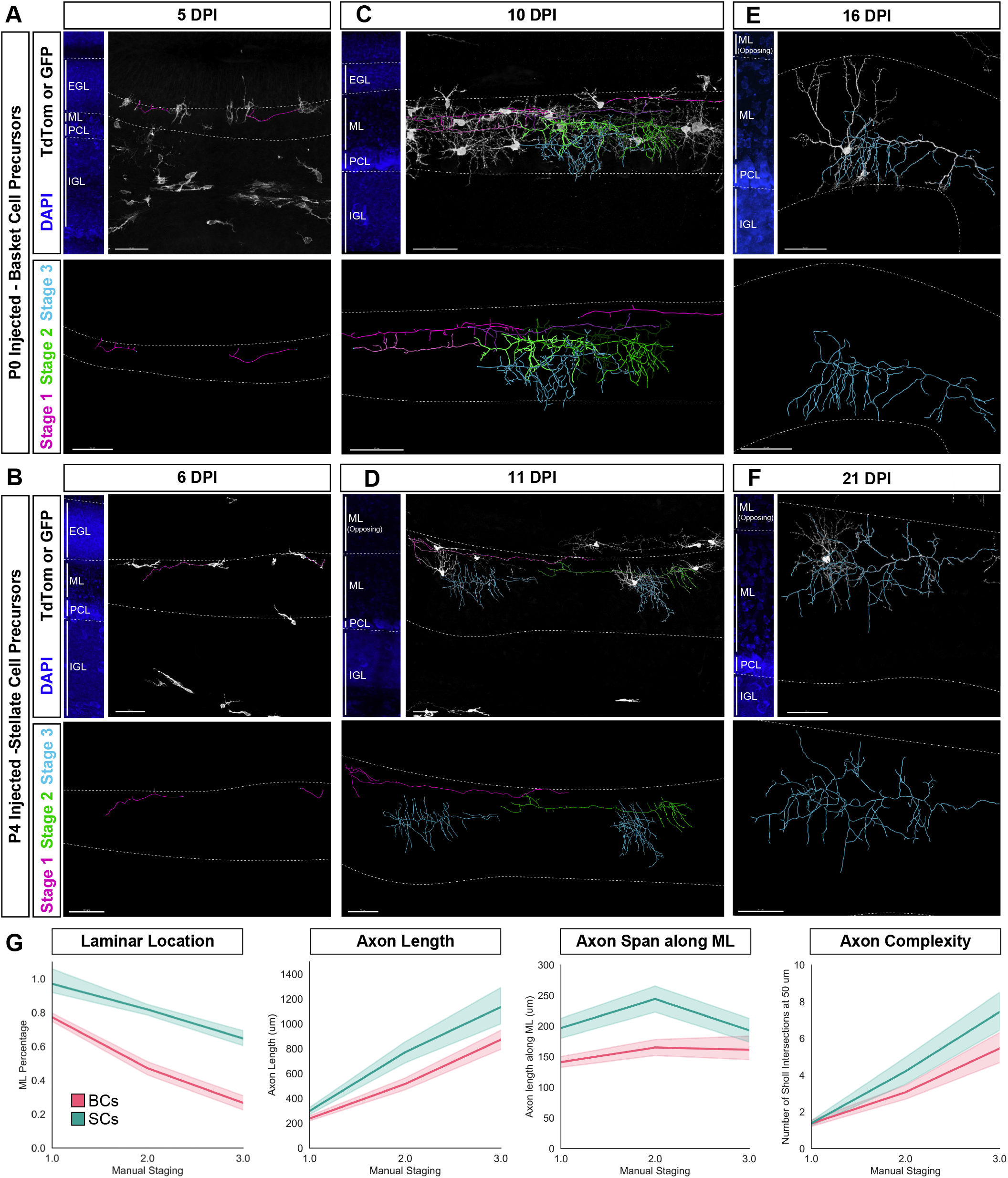
Comparisons of axonogenesis between BC- and SC-fated MLIs. **(A-F)** Fluorescently-labeled MLIs (*top panels,* greyscale) and axonal reconstructions (*top and bottom panels).* BCs were labeled by TMX injection at P0 **(A,C,E)** and SCs were labeled by P4/5 injection **(B,D,F)**. Dashed lines demarcate the ML. Images of DAPI-stained cerebellar cortex are shown in the *top left* panels. **(A)** Migrating BCs at 5 days post-TMX injection (DPI) extend a trailing process, the presumptive axon (magenta). **(B)** Migrating SCs at 6 DPI are in the superficial ML and extend a simple trailing process (magenta). **(C)** At 10 DPI, BCs display different stages of axonogenesis and are stratified in the ML according to axonal arbor progression. Immature BCs with a simple axon are located at the apical ML (pink or magenta traces, consistent with manual stage 1 of maturation). BCs with increasing axonal branching are located in the middle ML (green traces, Stage 2 MLIs; and blue traces, Stage 3). **(D)** SCs at 11 DPI are similarly stratified in the ML according to progression of axonogenesis (Stage 1, magenta; Stage 2, green; Stage 3, blue). Note that the axonal spans of SCs at this stage are considerably larger than BCs at similar stages. **(E)** Representative BC axonal morphology nearing maturity (16 DPI, blue). **(F)** Representative SC axonal morphology nearing maturity (25 DPI, blue). All scale bars are 50μ m. **(G)** Line plots of single morphological parameters along stages of maturation, as per manual staging. BCs and SCs differ significantly at all stages of axon morphogenesis for ML laminar locations occupied, axon length, axon span along the ML, and axon complexity.

In this visual analysis, stage 3 BC and SC subtypes are distinguishable based on differences in arbor geometry, laminar location, and axonal targeting. Quantifications of individual parameters further confirmed our pseudotime findings that BCs and SCs segregate throughout axon morphogenesis (Figure 8G). However, subtype differences were not visually apparent to the trained eye at stage 1 and to a large extent at stage 2, as features common to maturing interneurons visually outweigh minute subtype-specific differences. The inability to resolve the early segregation by eye further emphasizes the power of large-scale, multi-dimensional analyses. Taken together, visual inspections of MLI morphogenesis within the native ML environment confirmed pseudotime outputs.

### A proof of concept for morphology-based predictive cell-typing of single neurons

So far, we have shown that subtyping MLIs can be achieved at the population level using pseudotime trajectory inference. We next asked whether our dataset and approach may be useful for subtyping novel MLIs with single neuron resolution. We reasoned that by embedding single MLI traces into the established PHATE-generated trajectory, one could infer characteristics such as maturity and identity, using analogous properties of the nearest residing neighbor. As such, we propose the terms ‘pseudo-maturity’ and ‘pseudo-identity’, as the two cells likely do not reside within identical locations within the multi-dimensional data space, but one will nonetheless inform the identity of the other (Figure 9A). As a proof of concept, we saved the coordinates of the PHATE pseudotime manifold as a baseline operator (Figure 9B), and obtained new MLI reconstructions and their morphometric signatures (Figure 9C). The novel cells were embedded into the PHATE manifold as a new data point, followed by identification of the nearest neighbor (Figure 9D). We confirmed that the ‘pseudo-identity’ for both of the cells analyzed matched that of their true terminal identity, as determined by the corresponding time of TMX induction (Figure 9D). Given results from the nearest neighbors analyses, we expect the accuracy of predicted ‘pseudo-identities’ to be within the 75-90% range (Figure 7G). Taken together, we present a resource for current and future MLI researchers for predictive morphological cell-typing, and offer a proof of concept for other neuronal populations.

**Figure 9.**
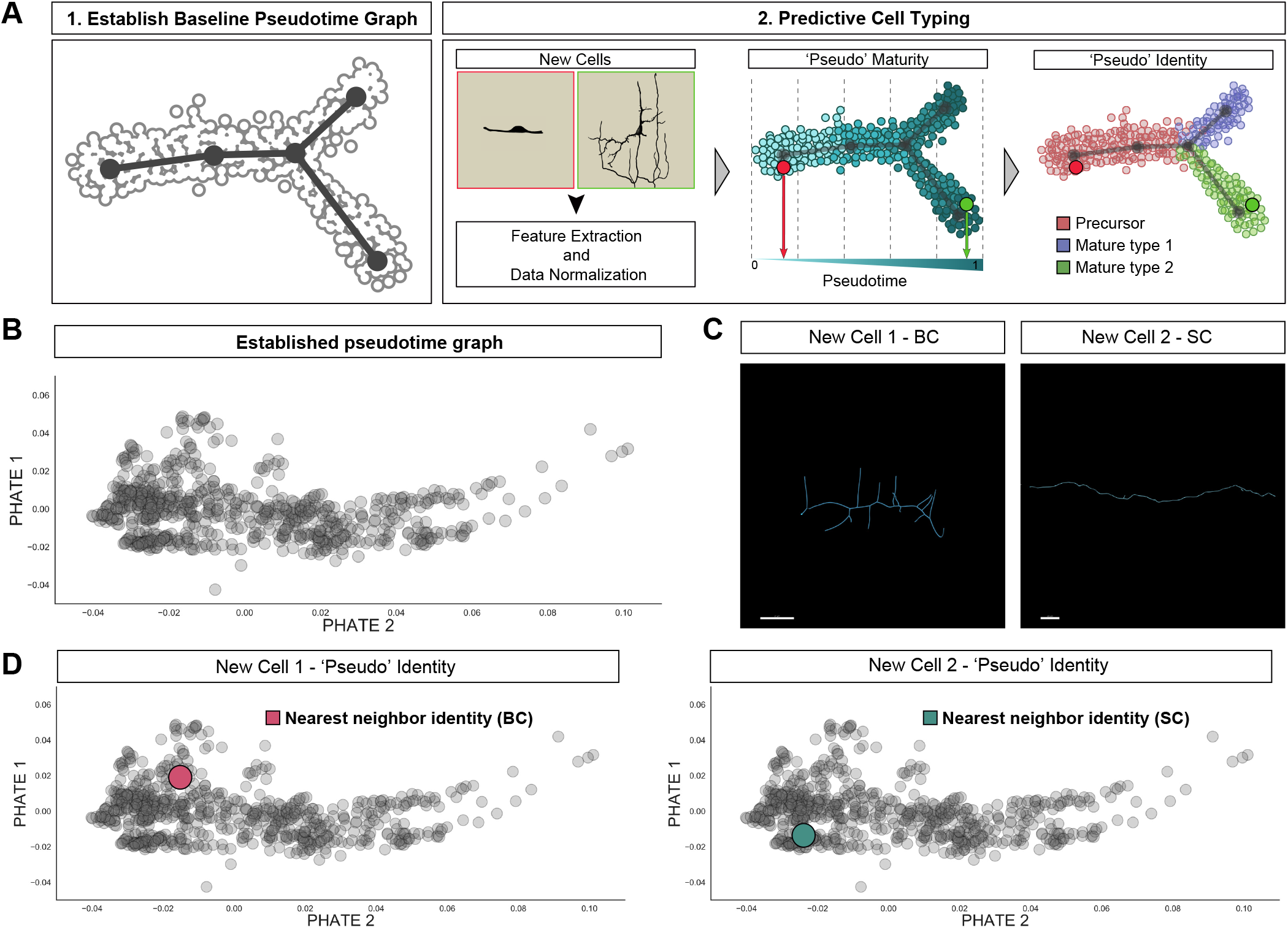
Proof of concept for predictive cell typing using morphological pseudotime. **(A)** Schematic for workflow of predictive cell typing using morphometric features. 1. Establish baseline pseudotime framework through large-scale single neuron profiling. 2. Obtain reconstructions for new cells, using previously established parameters. 3. Normalize new cell data to dataset. 4. Determine “pseudo-maturity” (*left*) pseudo-identity *(right)* of new cells. **(B)** Pseudotime trajectory established from Figure 7. **(C)** Reconstructions for two new cells that were not included in previous pseudotime analyses. Cell 1 was labelled at P0, and therefore is a presumptive BC, while cell 2 was labelled at P5, and therefore is a presumptive SC. **(D)** ‘Pseudo’ identities of new cell 1 (*left*) and new cell 2 *(right).* Color of highlight indicates the identity of the nearest neighbor (BC cell 1 and SC for cell 2). Scale bars are 20μm.

## Discussion

Over 130 years ago, Santiago Ramón y Cajal took advantage of the accessible and compact organization of the cerebellar MLIs to propose and substantiate the neuron doctrine (Cajal 1888; Cajal 1911; Sotelo 2015). In homage to the original master, we re-examined the morphological diversity of MLIs using genetic and computational methods of the modern era. We show through clustering and dimensionality reduction approaches that MLIs divide into the basket and stellate cell subtypes, thus demonstrating that MLI heterogeneity is best described by the two populations rather than the continuous one population model. Iterative removal of morphological parameters identified the axonal arbor as the defining feature for MLI classification. Building on this finding, we compared axonogenesis of basket- and stellate-fated cells by combining genetic lineage tracing with a novel pseudotime trajectory inference approach. To validate this approach, MLI reconstructions were spatially and temporally annotated and visualized to confirm their developmental progression along pseudotime. In doing so, we determined that MLI subtype identities emerge during migration prior to reaching sites of final integration. Finally, we present our dataset as a tool for predictive subtyping of MLIs and offer a proof-of-concept for other neuronal populations.

Our study demonstrates the power of large-scale morphological quantifications to define neuronal subtypes and the anatomical and developmental features that distinguish them. As this approach can be widely applied to study morphological diversification, we highlight strengths and limitations. First, we identified a BC-SC division at maturity supported by multiple algorithms. Since we adhered to conservative clustering rules, it is possible that the heterogeneity among SCs may parse into further subtypes with additional modalities. For instance, the early-born SCs in the lower ML presented divergent migratory and morphological features during pseudotime compared to the remaining SCs, suggesting that this population might represent another discrete subtype. Second, genetic methods for targeted and high-resolution labelling of MLIs allowed for sampling throughout their developmental progression. Large numbers of traces were required to adequately capture all stages of MLI morphogenesis, which can slow throughput due to the need for sparse labelling. Sampling can be improved by multicolor fluorescent reporters. Third, we provide the first demonstration of pseudotime trajectory inference using neuronal morphological information. As such, our analyses were dependent on pseudotime and modeling algorithms developed for transcriptomic and proteomic datasets. While we obtained a reasonably robust trajectory for aligning MLI identities, this approach might benefit from algorithms optimized for morphology. Fourth, lineage tracing and pseudotime ordering of snapshots acquired across two weeks of development enabled the characterization of MLI morphogenesis. Although morphological pseudotime is an approximation of changes occuring in real time, it proved useful for delineating subtype-specific development and extracting statistically meaningful features. Altogether, the application of trajectory inference methods to quantitative morphological data is not only novel but also produced new insights into MLI diversification, which we detail below.

### A revised taxonomy of MLIs

The long-standing debate of whether MLIs constitute one or multiple cell types reflects an enduring motivation to classify neurons into a systematic framework. The classical basket/stellate cell division is based on canonical patterns displayed by deep BCs and superficial SCs, which in our study only accounted for half of the total MLI population (Eccles et al. 1967; Palay and Chan-Palay 1974; Sotelo 2015). From observations of intermediate BC-SC morphologies, Cajal proposed that MLIs constitute a single cell type with continuously varying properties (Cajal et al. 1995; Sotelo 2015). In support of the ‘one population’ model, several studies demonstrated systematic variations of anatomical and electrophysiological features across molecular layer depth (Paula-Barbosa et al. 1983; Rakic 1972; Rieubland et al. 2014). Multivariate analysis of MLI anatomy similar to the one taken here failed to segregate the population by principal component analysis (Sultan and Bower 1998). This study was limited to twenty-six MLIs and contained few cells in the upper molecular layer due to the stochastic nature of Golgi labelling. With genetic access to MLIs that enhanced sampling across laminar positions, we obtained a clear BC-SC division among our dataset, which was confirmed through UMAP, t-SNE, hierarchical clustering, and statistical analyses.

Interestingly, the continuous heterogeneity we observed across the SC group reconciles both sides of the MLI taxonomy debate. Several SC morphometric quantifications, including dendritic and soma features, co-varied with laminar position in a manner that is consistent with the continuous variation previously attributed to the entire MLI population (Cajal et al. 1995; Rakic 1972; Rieubland et al. 2014; Sultan and Bower 1998). Moreover, we did not observe bimodal distributions for any of the morphological parameters that would indicate further divisions. Together, our findings support a continuous variation model for describing the SC population. The spatial variation in SC identity is not limited to anatomical properties but extends to functional ones. Variations in MLI connectivity patterns across the molecular layer depth exert different influences on MLI network activity and Purkinje cell firing (Arlt and Häusser 2020; Rieubland et al. 2014). Increasingly, single cell data are revealing the prevalence of continuous variation in the organization of neuronal populations (Cembrowski and Menon 2018; Cembrowski and Spruston 2019). For instance in the striatum and hippocampus, graded RNA expression within discrete cell types also correlate with spatial positions that reflect local or gross (i.e. dorsal-ventral axis) anatomical organization (Gokce et al. 2016; Arlt and Häusser 2020; Muñoz-Manchado et al. 2018; Harris et al. 2018; Cembrowski et al. 2018; Cembrowski et al. 2016). A few examples of neuronal subtypes that exhibit continuous variation in molecular and electrophysiological signatures have been highlighted so far (Muñoz-Manchado et al. 2018; Gouwens et al. 2019; Scala et al. 2020), but meaningful relationships between variations in molecular, functional, and morphological properties remain unknown. Single cell characterizations of transcriptional and functional properties of cerebellar neurons are underway (Zeisel et al. 2018; Kozareva et al. 2020). A further understanding of MLI biology will come from investigations of correspondence between modalities.

### MLI axonogenesis begins during migration

In this study, we demonstrated the coupling of MLI axonogenesis with migration. Live imaging studies of MLI migration have reported the extended, circular trajectory that includes four distinct phases of radial and tangential migration through the molecular layer depth (Cameron et al. 2009; Wefers et al. 2017; Wefers et al. 2018). These studies did not distinguish between basket and stellate cells. Through pseudotime trajectory inference and confirmation in situ, we show that MLI axon development is prolonged and temporally overlaps with multiple phases of migration. Both young BCs and SCs in the apical ML exhibit elongated soma morphologies and extend a single prospective axon. The timing, location, and migratory morphology of these MLIs are consistent with the first tangential phase of ML migration (Cameron et al. 2009). These simple neurites are consistent with descriptions of trailing processes that emerge during migration of some interneuron populations (Lim et al. 2018; Marín et al. 2010; Sakakibara and Hatanaka 2015). In the cortical plate for example, a subpopulation of radially migrating interneurons leave behind a trailing process that remains stationary and eventually develops into the axonal arbor (Lim et al. 2018). Live imaging studies of MLI migration did not report the presence of trailing processes nor axon formation (Wefers et al. 2017; Cameron et al. 2009). Pseudotime modeling on the other hand, indicates that MLI axonal arborization initiates with a single trailing process that elongates along the apical ML prior to the emergence of collaterals and branch remodeling. In a study which charted BC axon morphogenesis, secondary axon collaterals extended towards the PC somas through guidance from extracellular guidance cues, as opposed to direct cell-cell interactions (Telley et al. 2016). Although they did not account for the displacement of BCs during the process of axonogenesis, our findings are consistent with the existence of such extracellular gradients for directing MLI migration and axon guidance. Future studies on MLI migration should shed light on the identity and properties of these molecules, which will likely resemble a gradient along the ML.

### Early emergence of MLI identities

MLIs are the last category of interneurons generated in the cerebellum, as part of the Pax2-expressing GABAergic interneuron lineage that originates from a single multipotent pool of progenitors in the ventricular zone (Hoshino et al. 2005; Leto et al. 2006; Zhang and Goldman 1996; Maricich and Herrup 1999; Weisheit et al. 2006). Proliferating MLI precursors exit the ventricular zone and migrate to their secondary germinative region in the cerebellum, the prospective white matter (PWM; Leto et al. 2006; Sotelo 2015). As such, MLIs represent an interesting example in the nervous system where proliferating progenitors migrate to an intermediate zone for further expansion and terminal division. Since MLI fates are not intrinsically determined nor restricted at birth, they are influenced by instructive cues encountered as they migrate from the PWM to their final position in the molecular layer (Leto et al. 2009). Intriguingly, birth-dating studies have demonstrated that post-mitotic MLIs remain in the PWM for 1-4 days following terminal division (Leto et al. 2009). In Leto et al., comparisons between transplantation of dissociated neurons and solid PWM grafts suggested that the PWM microenvironment may be a source of instructive cues for MLI fate specification. The significance of this protracted and variable period of development in the PWM on MLI differentiation remains unclear. Moreover, whether the inside-out placement of MLIs and subtype-specific phenotypes within the mature ML is restricted by birthdate or the time of PWM exit is unknown.

Through genetic lineage tracing and pseudotime trajectory inference, we determined that MLI subtype identities emerge during early phases of migration in the ML, and that a subset of SCs are generated along with BCs. The BC and SC lineages were separable across developmental stages for a number of morphometric parameters, including total axon length, axon arbor span, and laminar locations. This early divergence was further confirmed by the Palantir- and PHATE-generated pseudotime trajectories, together revealing cluster-specific enrichments for BC- or SC-fated cells along all developmental states. Although transplantation studies suggest that commitment to MLI terminal identities remains plastic (Leto et al. 2009), our data indicate that MLI fates are instructed earlier. Moreover, we demonstrated that MLI subtype identities are not dictated by ML laminations. Lower ML SCs are among the early-born MLIs, occupy similar laminar locations as BCs, but were separable from the BC lineage through pseudotime. Additionally, SCs are not restricted to upper ML positions but instead are distributed through the depth of the ML. Together, we propose that MLI laminar positioning is determined at the time of terminal division. However, subtype identities are shaped by the timing of PWM exit or cues from the nascent ML. Although BCs and lower ML SCs are born at similar times, lower ML SCs might adopt a different fate by taking longer to traverse through the PWM and ML prior to axonogenesis. MLI lineage-specific profiling and manipulations are required to further identify the source and timing of subtype specification.

### Concluding Remarks

In conclusion, our study leveraged computational methods to characterize MLI development with single-cell resolution and temporal coverage, while annotating individual cells using anatomical and lineage information. Such approaches will inform future studies for cues that shape neuronal patterning and for linking local variations in connectivity patterns to function. Importantly, restoring anatomy and spatial-temporal contexts to single-cell profiling will provide the integrated descriptions needed for understanding how diverse and complex neuronal subtypes arise and assemble into circuits.

## Material and methods

### Mouse strains

Mouse lines used in this study have previously been described. GABAergic neuron targeting *Gad2-ires-Cre* (JAX accession #: 010802; Taniguchi et al. 2011) and MLI-specific *Ascl1-CreERT2* (JAX #: 012882; Kim et al. 2011) were obtained from Jackson Laboratories. Cre reporter lines Ai14 *Rosa-Cag-LSL-TdTomato* (JAX #: 007908; Madisen et al. 2010) and Rosa^mT/mG^ (JAX #: 007576; Muzumdar et al. 2007) were obtained from Jackson Laboratories. Mice were maintained on a C57/B6J or mixed C57/B6J and FVB background.

All experiments were carried out in accordance with the Canadian Council on Animal Care guidelines for use of animal in research and laboratory animal care under protocols approved by the Centre for Phenogenomics Animal Care Committee (Toronto, Canada) and the Laboratory Animal Services Animal Care Committee at the Hospital for Sick Children (Toronto, Canada).

### Virus labelling of neurons

Recombinant Brainbow AAV9-hEF1a-LoxP-TagBFP-LoxP-eYFP-LoxP-WPRE-hGH-InvBYF and AAV9-hEF1a-LoxP-mCherry-LoxP-mTFP-LoxP-WPRE-hGH-InvCheTF viruses (Cai et al. 2013) were obtained from Pennsylvania Vector Core. ~1 x 10^12^ viral genome particles per mL of each Brainbow virus was prepared in sterile phosphate-buffered saline (PBS, pH = 7.4).

To introduce virus into the cerebellum, P0 - 14 pups (P0 for PCs, P5-14 for MLIs) were anaesthetized with either ice (P5 or younger) or isoflurane using a rodent anesthesia machine (>P6 and older, 4% in O2). A 25G needle was used to make a small puncture into the caudal-medial position of the right cortical lobe, and 1.5 uL of rAAV virus was injected into the lateral ventricles with a Hamilton syringe and 33G blunt-ended needle. Injection procedures were repeated for the left cortical lobe for bilateral labeling of neurons. Animals were sacrificed and cerebellums dissected at P75 for mature classifications.

### Tamoxifen induced labelling of neurons

Tamoxifen (Sigma, T5648) was dissolved in corn oil (Sigma, C8267) to a concentration of 10 mg/mL (high dose injections; Figure 5C, *middle and right* panels), 0.5 mg/mL (low dose injections for sparse labeling using the mTmG Cre reporter), or 0.05 mg/mL (low dose injections for sparse labeling using Ai14-TdTomato Cre reporter). P0 - P7 pups were put on ice for 1 minute for anesthesia and to reduce oil leakage following injections. A 0.5cc insulin syringe (BD) was used to introduce 10 - 20 uL of tamoxifen solution through intraperitoneal (I.P.) injections. The needle tip was held inside the pup for 20 seconds to prevent excessive leakage. Subcutaneous injections can be substituted for labeling of stellate cells (P4 - P7 injections), but I. P. injections are necessary for capturing basket cells in our postnatal injection scheme due to its faster acting nature for tamoxifen introduction and activation. Pups were sacrificed at least 3 days post-injection.

### Histology

Mice were either anesthetized by hypothermia (P5 or younger) or under isofluorane (4% in O2 for induction; 1.5-2% in O2 for maintenance), and transcardially perfused with physiological saline solution (0.9%; Baxter) followed by 4% paraformaldehyde (PFA) in PBS. Animals were perfused by the gravity perfusion method. Brains were post-fixed in 4% PFA overnight at 4°C or for 3.5 hours at room temperature (RT).

100 um sagittal sections of the vermis cerebellum were sectioned using a vibratome (Leica). Sections were incubated for 3.5 hours in blocking buffer (0.5% Triton-X, 4% normal donkey serum in PBS), and incubated for 72 hours at 4°C with primary antibodies. Following 3 x 15 min PBST (0.5% Triton-X) washes, sections were incubated for 3.5 hours at RT with Alexa-conjugated secondary antibodies (Invitrogen or Jackson ImmunoResearch). Sections were mounted onto glass microscope slides, coverslipped using Fluoromount G (Southern Biotech). Nuclei were labelled using DAPI or NeuroTrace Nissl 435/455 (Invitrogen).

Primary antibodies used for this study were as follows: chicken anti-GFP (1:2000, Aves Laboratories, GFP-1010); rabbit anti-mCherry (1:500, Kerafast, EMU106); rat anti-TFP (1:500, Kerafast, EMU104), guinea pig anti-TagBFP (1:500, Kerafast, EMU108); Rabbit anti-RFP (1:1000, Rockland, 600-401-379); goat anti-Parvalbumin (1:1000, Swant, PVG213), and rabbit anti-Calbindin (1:1000, Sigma, C9848).

### Confocal imaging

Images of single MLIs were taken on a Leica SP8 scanning confocal microscope, using a 40X oil objective (NA = 1.3). Z-stacks were collected with a 0.5 um (mature dataset) or 1 um (developmental dataset) step size throughout the depth of the cells, to encapsulate the entire dendritic and axonal arbor. Only neurons where arbors were not cut off by the vibratome were imaged and analyzed. The Z-compensation feature was used to avoid signal saturation throughout the depth of the tissue. Images were acquired with slightly different XY pixel sizes to accommodate varying arbor sizes, and to be able to capture single MLIs within one field of view. XY pixel sizes used were maintained at ~135 nm.

### MLI counts

To account for the percent laminar distribution of singly labelled MLIs following Brainbow injections, we divided the ML into four strata, and counted the number of labelled MLIs within each strata. The XScope app was used to create a 1 x 4 grid, which was resized to match the area of the ML. The number of MLI were then counted within each strata, using the cell counter plugin in FIJI (Schindelin et al. 2012). Each counting frame additionally consisted of one inclusion and one exclusion edge, and cells were counted if found entirely within the counting frame or overlapping with the inclusion edge but not the exclusion edge.

### Manual curation of MLI morphologies

Canonical BCs and SCs were identified according to standards in literature (Amat et al. 2017; Sergaki et al. 2017; Gaffield and Christie 2017; Sultan and Bower 1998). Canonical BCs were identified by: 1) soma location within the bottom ⅓ of the ML, 2) a long horizontal axonal shaft which gives rise to basket terminals, and 3) a dendritic arbor which reaches the apical ML. Canonical SCs were identified by: 1) soma location within the upper ½ of the ML, 2) absence of basket terminals, 3) radially oriented dendritic and axonal arbors. The manually curated set of BCs likely contained fewer cells than both hierarchical clustering and UMAP analyses due to our strict laminar location cut-off.

### Morphological reconstructions

3D reconstructions for single neuron morphologies were completed in Imaris (Bitplane). Dendritic and axonal arbors were semi-automatically reconstructed using the filament tracer (autodepth mode). Surface render of the cell soma was performed using the surfaces module. We note that due to the membrane targeted nature of the Brainbow fluorophores, the soma volume data for mature MLIs more accurately describes the volume of the cell membrane, and not the entire somata.

### Data compilation of single neuron anatomical features

For each neuron, we: 1) registered the spatial coordinates to note the cell location and folia of origin in the vermis cerebellum; 2) performed surface rendering for the cell soma, 3) performed morphological reconstructions, and 4) extracted 27 quantitative features to describe the dendrites, axon, somata, and location (Table 1).

Basic morphological features were exported from Imaris using the statistics tab. These include: axon/dendrite length, number of Sholl intersections at 10, 50, 100, 150, and 200 um, mean/max branch level, axon straightness, and volume of somatic render. The remaining features were manually compiled:

#### Mature dataset

1. The number of filopodia was calculated as the number of terminal branches under 1.5um in length.
2. Filopodia density was calculated by normalizing total filopodia numbers to the dendrite length.
3. Z-depth, height of ML covered by dendrites/axons, and axonal span along the ML were measured using the measurement tool in Imaris.
4. Radial symmetry of a cell was manually graded between 1-4, but was not included in the final clustering dataset due to the qualitative nature of the assessments.
5. For analyses of upwards- and downwards-oriented axon collaterals (numbers, length, and percentage), collaterals directed towards the apical or basal ML were highlighted in Imaris, and the corresponding data were compiled from the statistics tab. For collaterals which were accurately angled, only branches more than 30 degrees away from the main axon were included in the analysis.
6. Presence of axon-carrying dendrites were noted for cells where the axon initial segment originates from the basal dendrites, not the soma.
7. Folial location was noted in folia 1, 3, 5, 6, 7, 8, 9, and 10. This data was not included in the final clustering analyses, but used for confirmation that MLI identities are not biased by the cell’s folia of origin.
8. Relative molecular layer (ML) position was measured using the measurement tool within slice mode. ML heights were calculated as the distance from the top of the PCL to the top of the ML. MLI soma heights were calculated as the distance from the top of the PCL to the centre of the MLI soma. MLI laminar locations were calculated as MLI soma height / ML height.
9. For each cell, the number of full baskets (one which fully envelopes the corresponding PC soma) with pinceau formations were counted in addition to the number of full baskets without pinceau formations, and the number of half baskets (one which less than entirely envelops the PC soma). Number of baskets were then calculated using a weighted system, according to the proxied functional significance of each basket. In this way, full baskets with pinceau formations were given a weight of 1, full baskets without pinceau formations were given a weight of 0.75, and half baskets were given a weight of 0.5.
10. The number of primary dendrites were counted as the number of branches which arises from the soma.

#### Developmental dataset

1. Overlapping parameters were compiled following similar protocols as the mature dataset. These include axonal height, depth and span information, measurements of upwards- and downwards-oriented axon collaterals, laminar locations, and the number of primary dendrites.
2. The number of axonal branches which reached into the PCL were counted, to account for developmental changes as most basket-forming cells have not formed full basket formations at the timepoints analyzed.
3. Soma diameter for each cell was measured along three dimensions using the measurement tool within slice mode (X = direction along the PCL, Y = direction perpendicular to the PCL, and Z = direction along parallel fibers). The final somatic volume was approximated using the ellipsoid volume equation: V = 4/3π x X-diameter x Y-diameter x Z-diameter.

We note that although a full 3D volumetric render of the soma similar to what was done for the mature dataset would have been more accurate, the process was time and computationally intensive. Approximating the soma volume allowed for the large-scale nature of our study.

### Clustering analyses

Prior to morphometric clustering, each dataset was zero mean standardized as previously described (Lanjakornsiripan et al. 2018). t-SNE was performed with the Rtsne package (version 0.15) for the R statistical environment (R Foundation for Statistical Computing 2018; version 3.5.1) with the following parameters: seed = 510, theta = 0, max_iter = 10,000, and perplexity = 7. The elbow method and silhouette analysis were performed using the NBClust package (version 3.0) in R. UMAP analysis was performed in Python (Python Software Foundation n.d.; version 3.7.3) using the umap and sklearn packages, with the following parameters: min_neighbors = 7, min_dist = 0.1, metric = ‘euclidean’. Hierarchical clustering was performed using the SciPy.cluster.hierarchy package in Python, using the Ward’s method (Ward 1963). SC subclades were identified visually through curation of the corresponding MLI traces.

### Camera Lucida illustrations

Camera Lucida traces of dendritic morphologies were created using the Affinity Designer (Serif) app on an iPad (Apple). Max projection of the dendritic arbors were obtained in FIJI. The resulting image was inserted in Affinity Designer. Traces of the dendritic and somatic outlines were completed using an Apple pencil.

### Expert directed pseudotime based on dendritic maturity

Manual staging of MLI morphometric maturation was performed based on our 4 stage maturation scheme (Figure 6 - Figure Supplement 1). Additionally, migratory precursor cells in Figure 6M were identified as neurons which had a migratory leading process (similar to those found on stage 1 cells), but which did not possess an observable trailing process.

### Pseudotime trajectory inference using Palantir

We adapted the Palantir pseudotime algorithm to align our single cell snapshots in the developmental space (Setty et al. 2019): 1) The developmental dataset was zero mean standardized outside of Palantir. This step was performed individually for the total MLI dataset, and the early-born MLI subset; 2) the normalized datasets were imported into Palantir as a normalized dataframe; 3) principal component analysis was performed to determine the principal components; 4) Diffusion maps of the data were obtained to determine the low dimensional phenotypic manifold of the data, with n_components = 5; 5) t-SNE representation of the data was created in the embedded space; 6) MAGIC imputation (van Dijk et al. 2018) was used to map trends for each morphological parameter onto the t-SNE map; 7) starter cells were manually assigned following expert curation of the corresponding morphology traces; 8) clustering was performed using the Phenograph package and the Louvain method (Levine et al. 2015), with k = 20.

### Trajectory inference using PHATE

We adapted the PHATE algorithm for aligning our single cell snapshots in the developmental space (Moon et al. 2019): 1) The developmental dataset was zero mean standardized in the same way as our Palantir analyses; 2) the normalized datasets were imported into PHATE as a normalized dataframe; 3) the dataframe was used to generate a PHATE estimator object with the following parameters: knn = 3; 4) diffusion pseudotime was performed on the PHATE operator using Scanpy (Wolf et al. 2018); 5) expert directed maturity stages were projected onto the PHATE manifold as a color label.

### Nearest neighbor analyses

The nearest neighbor analysis were performed following PHATE-based pseudotime trajectory inference by: 1) binning the pseudo-timeline into 0.2 unit increments; 2) identifying the nearest neighbor for each MLI reconstruction within the PHATE manifold; 3) determining whether the terminal identity of the nearest neighbor is the same or different from the cell in question; 4) determining the percentage of cells within each pseudotime bin with a neighbors of the same terminal fate.

### Experimental design and statistical analysis

Morphological reconstructions for the 79 mature MLIs were compiled from 9 P75 mice across cerebellar folias. Developmental axonal reconstructions for the 732 MLIs were compiled from 32 mouse pups, which included the following injection to collection timelines: 1) P0 to P5; 2) P0 to P7; 3) P0 to P8; 4) P0 to P10; 5) P0 to P13; 6) P0 to P16; 7) P0 to P25; 8) P1 to P13; 9) P1 to P16; 10) P1 to P25; 11) P2 to P13; 12) P2 to P16; 13) P2 to P27; 14) P3 to P13; 15) P3 to P16; 16) P4 to P10; 17) P4 to P13; 18) P4 to P15; 19) P5 to P13; 20) P5 to P14; 21) P5 to P16; and 22) P7 to P14. Animals of either sex were analyzed. The early born population for pseudotime analyses contained all P0 injected samples, as well as a small number of mature P1 labelled cells which were confirmed as BCs due to the elaboration of soma-targeting axons. The late-born population contained all P4-P7 injected samples. Statistical analyses were performed using the GraphPad Prism software or the SciPy package in Python. Means of two groups were compared using the two-tailed student’s t test with the Mann-Whitney nonparametric test. Graphs were created using the Seaborn or Matplotlib packages in Python, or using GraphPad Prism.

## Acknowledgements

This work was supported by a Canada Research Chair (Tier 2), a Sloan Fellowship in Neuroscience, an NSERC Discovery grant, a CIHR Project grant, and funding from the Hospital for Sick Children (to J.L.L.). We thank Dr. Shreejoy Tripathy and members of the Lefebvre lab for helpful comments on this manuscript, and Dr. Manu Setty for discussions on morphological pseudotime. We thank Scott Gigante for computational assistance for binning of pseudo-timelines.

## Author contributions

W.X.W and J.L.L. conceptualized and designed the project. W.X.W. performed experiments and computational analyses. W.X.W. and J.L.L. wrote the manuscript.

---

**Supplementary Figure S1.**
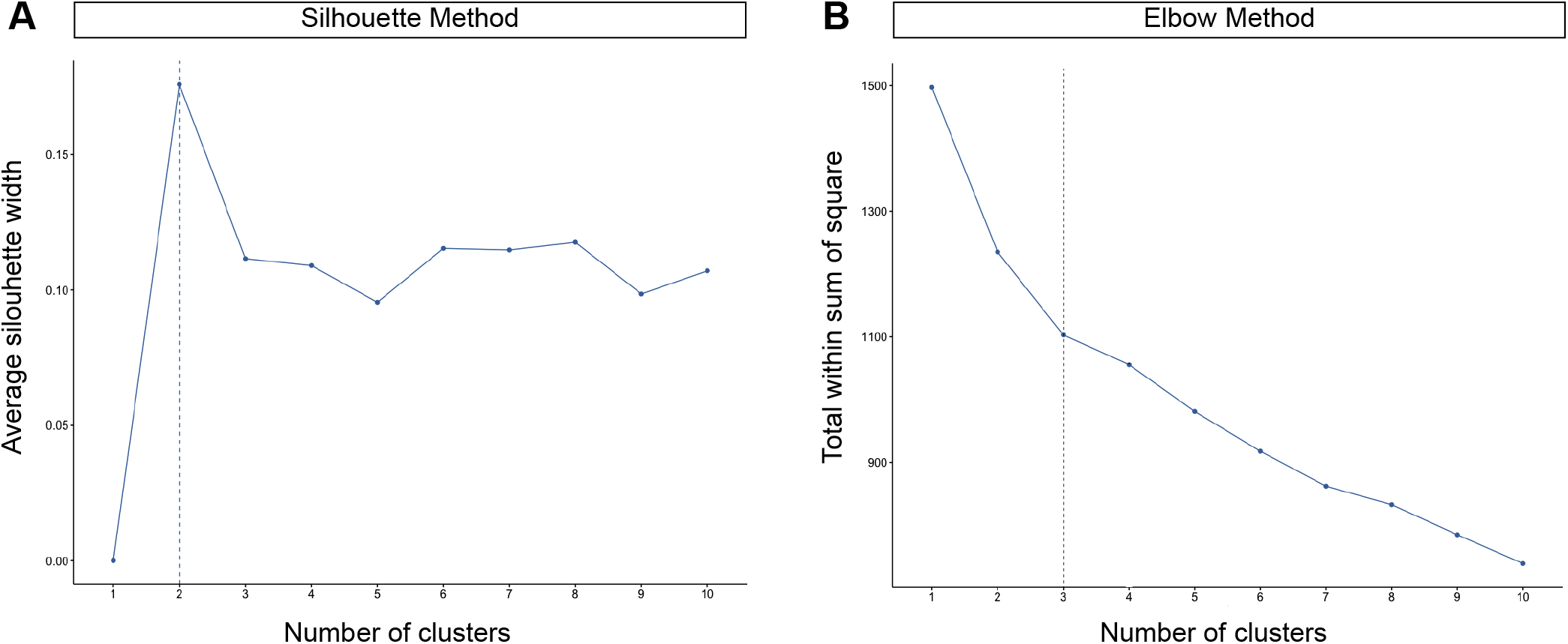
Statistical confirmation for the optimal number of MLI clusters. **(A)** The silouhette method measures the quality of clustering by assessing how well each cell lies within their cluster. **(B)** The elbow method measures the k-means score and the total within sum of squares distance by assessing the distance of each cell to the cluster centroid. The optimal number of clusters is determined as 2 by the silouette method and 3 by the elbow method.

**Supplementary Figure S2.**
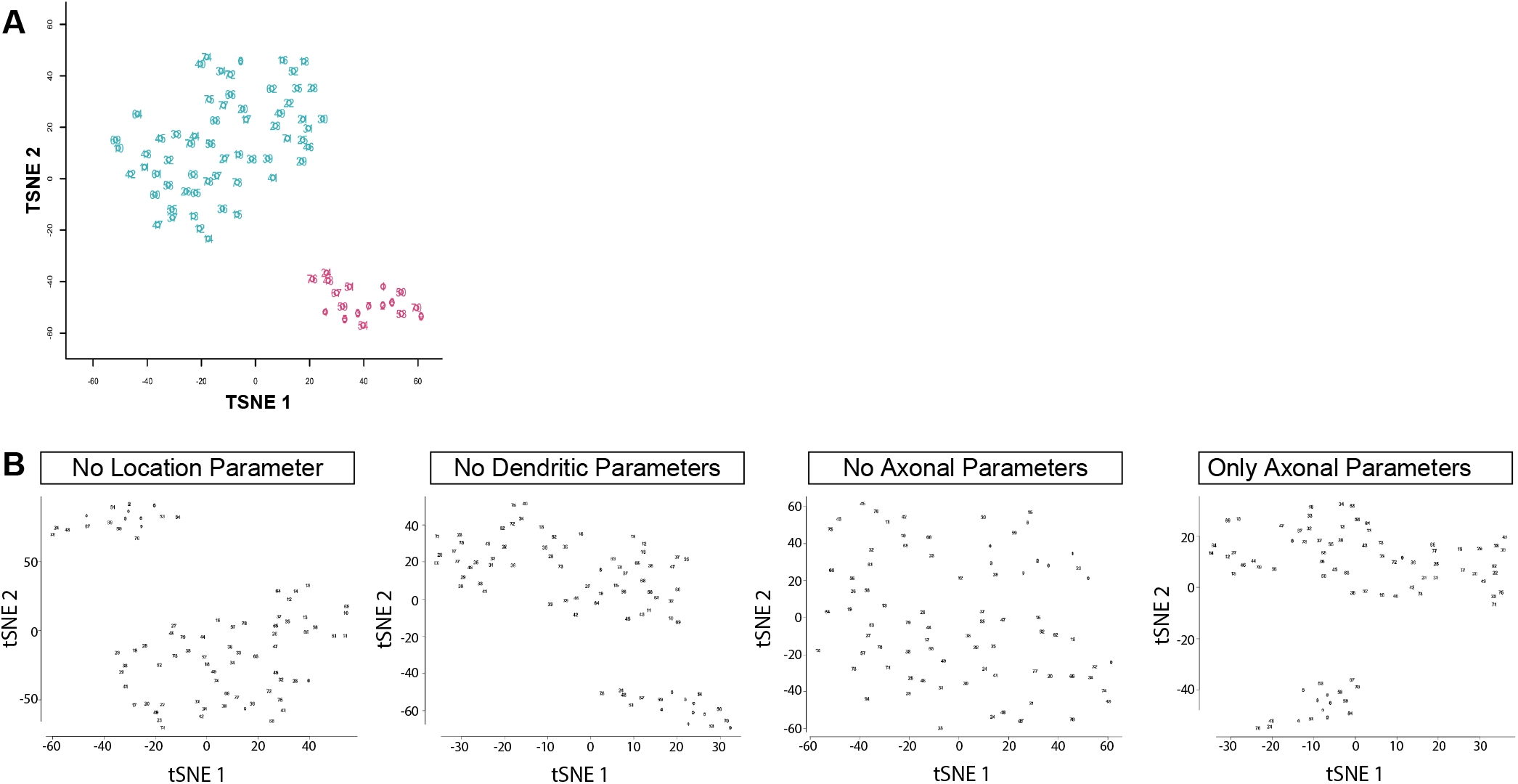
Confirmation of UMAP results by t-SNE. **(A)** t-SNE of 79 mature MLIs. BCs are clustered in pink and SCs are colored in cyan. **(B)** Iterative feature elimination using t-SNE confirms that axonal information is necessary and sufficient for the BC/SC division.

**Supplementary Figure S3.**
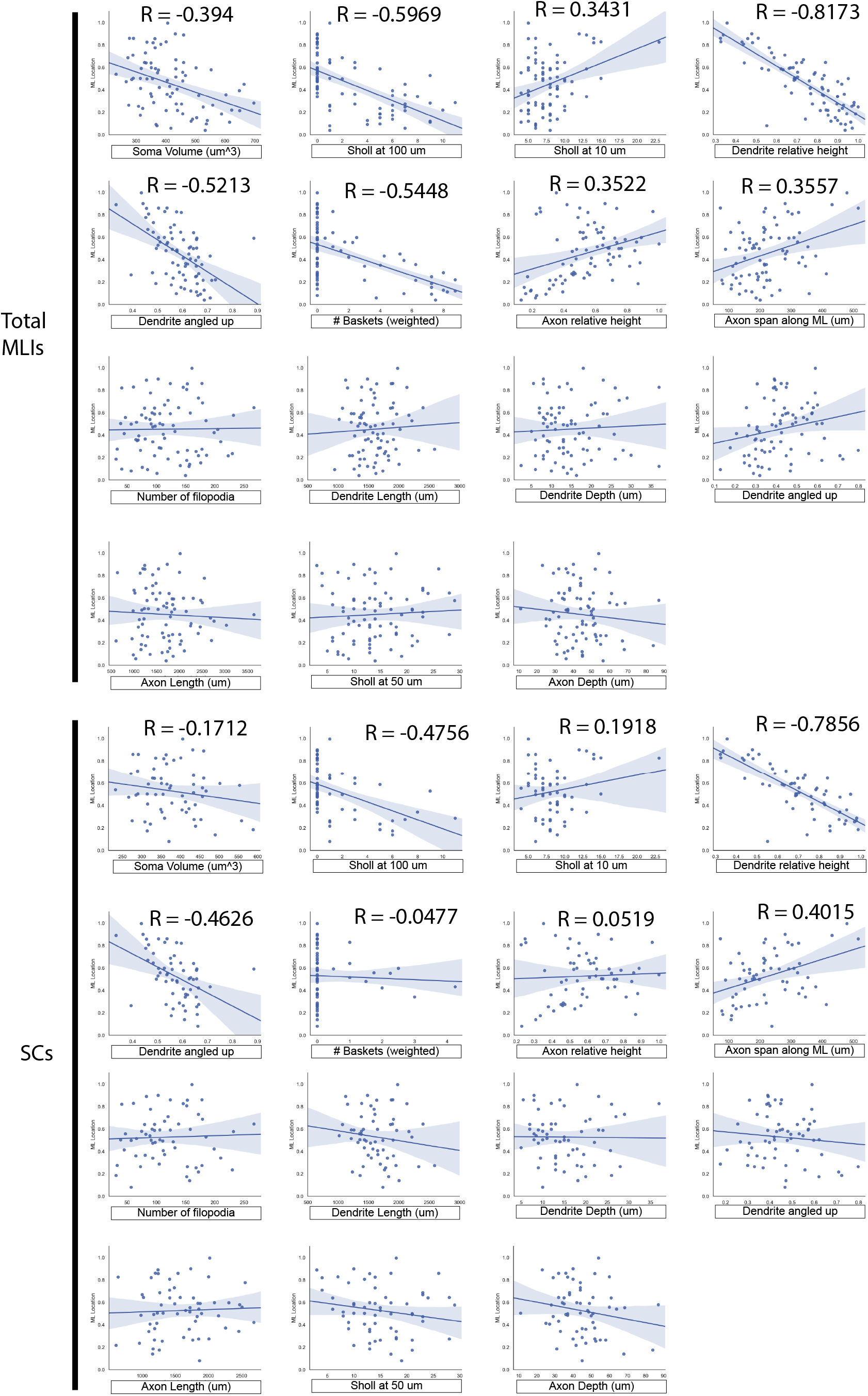
SCs present significant within-class heterogeneity as confirmed through linear regression of morphological parameters. The top eight parameters show a postive or negative correlation to ML depth for the entire MLI population.

**Supplementary Figure S4.**
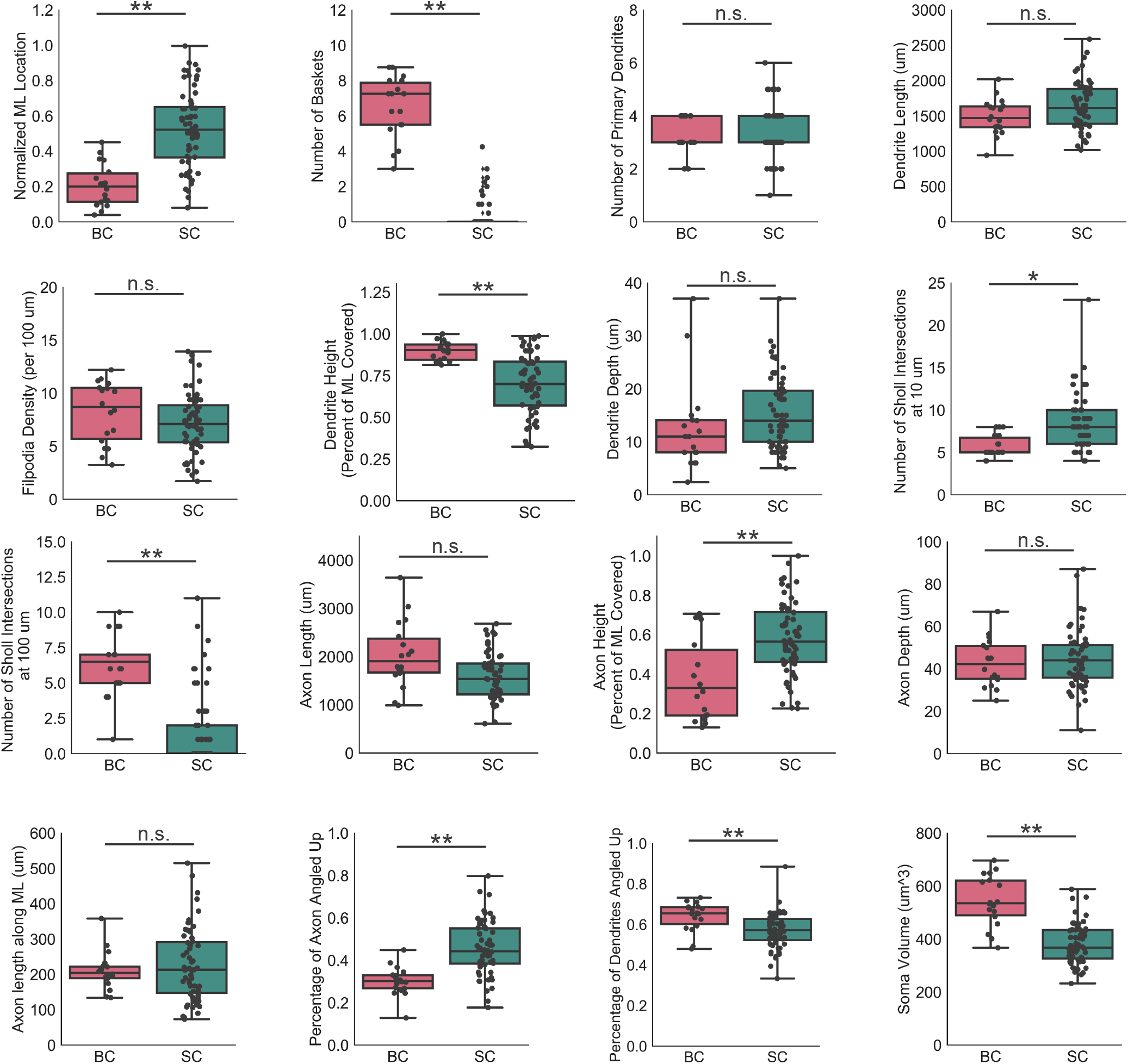
Individual morphometric parameters for BCs and SCs, as per hierarchical clustering. All parameters overlap between BCs and SCs, and thus no individual parameter is sufficient for MLI subtype identification. The number of baskets is a good proxy. All parameters follow a largely normal distribution, with no presence of bimodality. Upper and lower whiskers of the box plots represent the maximum and minimum values of the dataset, respectively. The box is drawn connecting the two innermost quartiles.

**Supplementary Figure S5.**
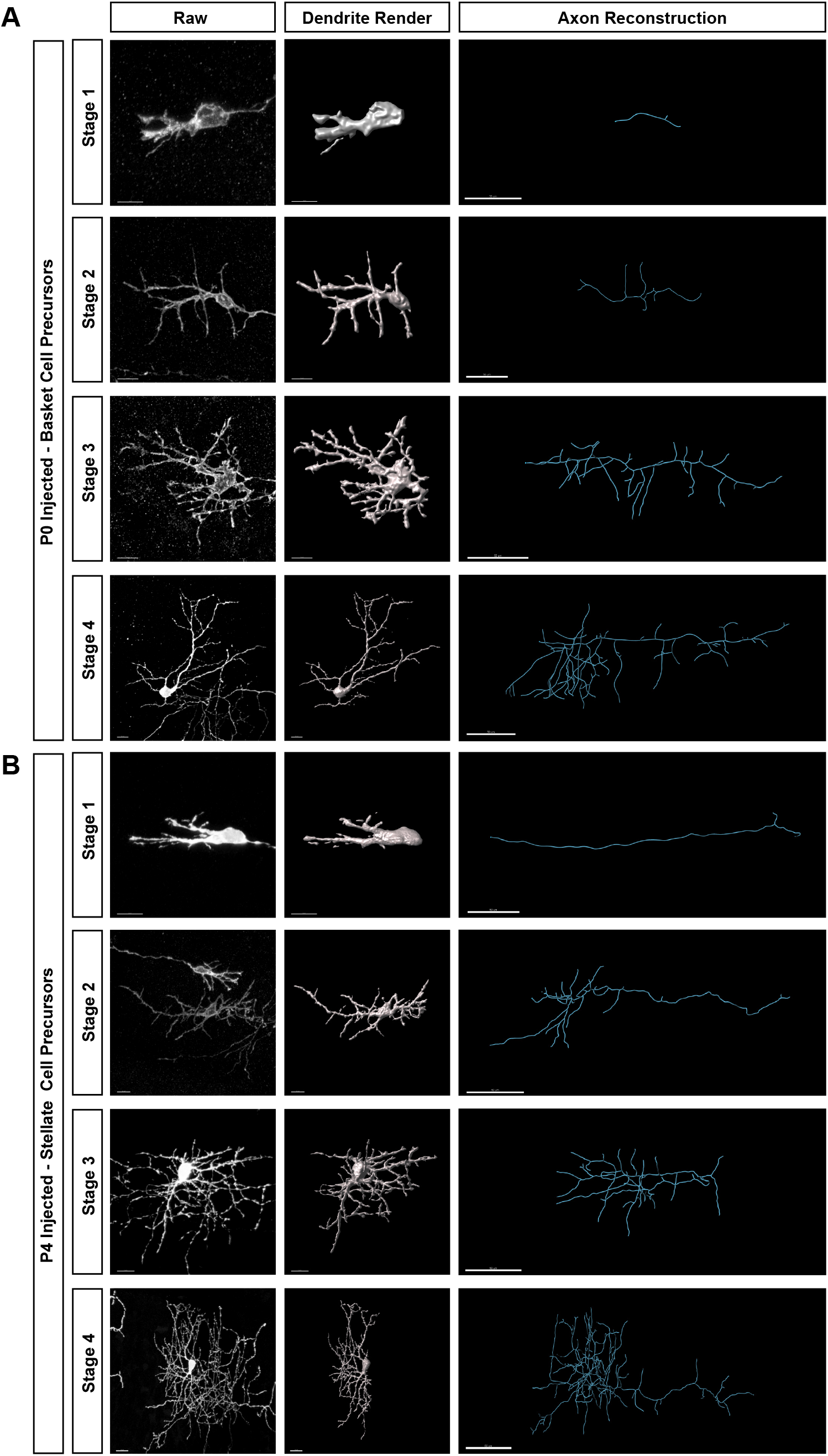
Manual staging of MLI morphogenic maturation based on dendritic morphology for **(A)** early (P0) and **(B)** late-born (P4-7) MLIs.Stage 1 cells are migratory MLIs with clear migratory morphologies and leading processes. Stage 2 cells are maturing migratory MLIs with a developing dendritic arbor. Stage 3 cells have dendritic arbors of increasing complexity and rounded somas. Stage 4 cells are mature MLIs with type-specific dendritic and axonal arbors.

**Supplementary Figure S6.**
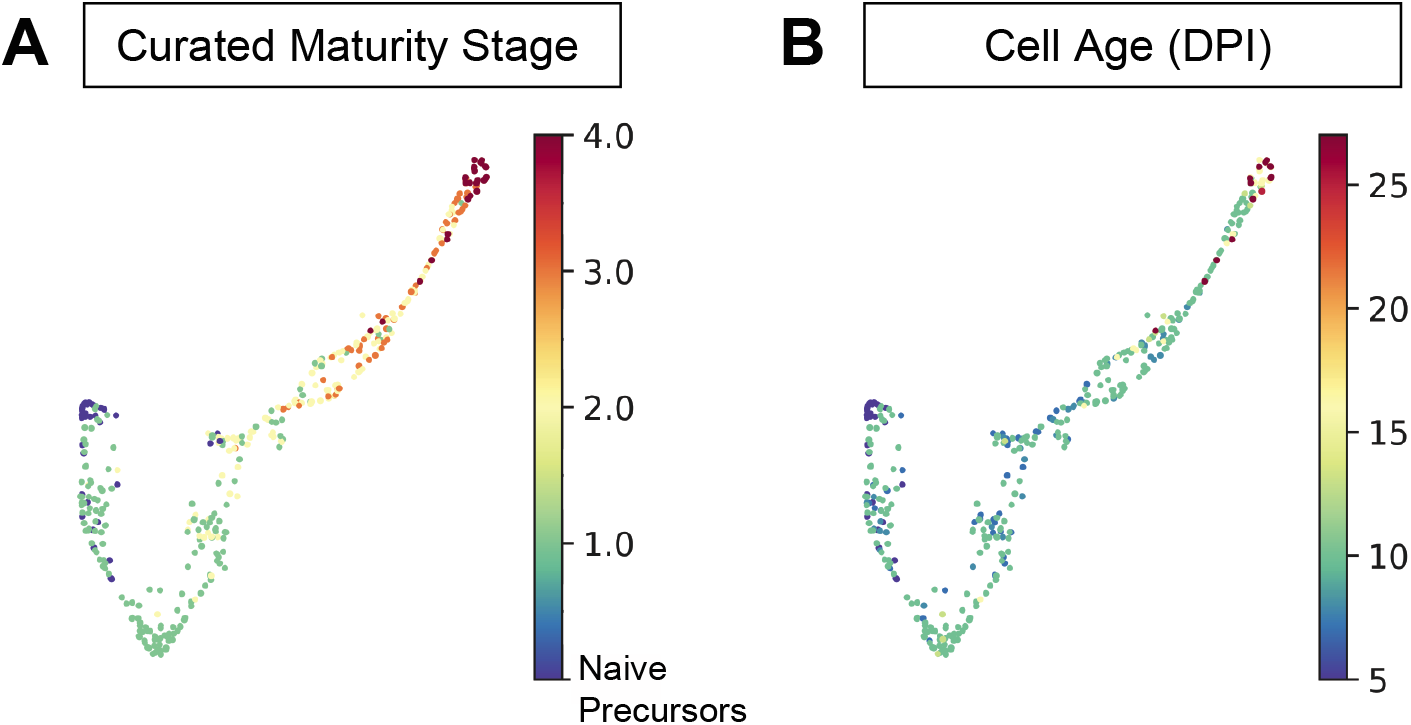
**(A)** Curated maturitystage for single MLIs projectedonto Palantir trajectory. Naive precursor cells correspond to figure 6M, and are characterized as migratory MLIs with prominent leading process, but lacking visible trailing process. **(B)** Cell age projected onto Palantir trajectory, as inferred from days after tamoxifen injection (DPI).

**Supplementary Figure S7.**
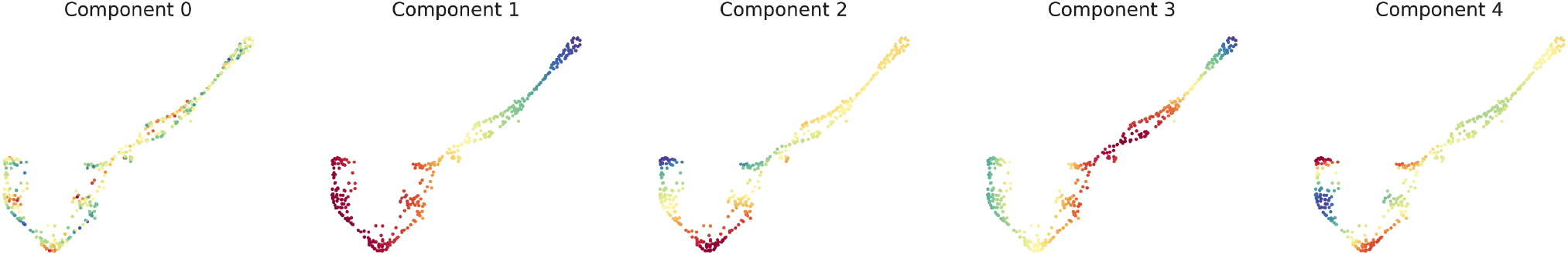
Diffusion map components for early-born MLI Palantir trajectory.

**Supplementary Figure S8.**
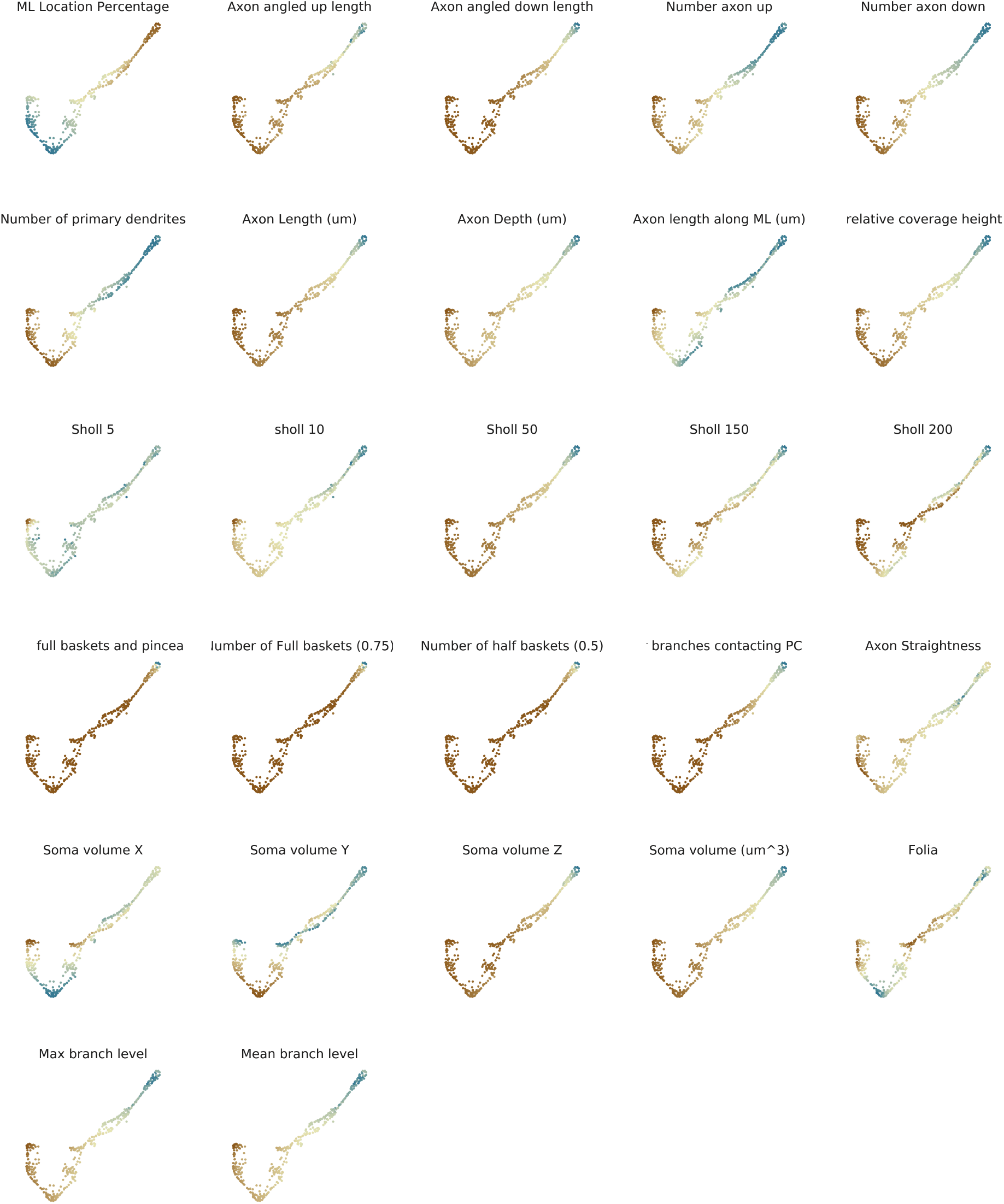
Projection of single morphometric parameters onto Palantir pseudotime trajectory. Minimum parameter values are highlighted in brown and maximum parameter values are highlighted in blue.

**Supplementary Figure S9.**
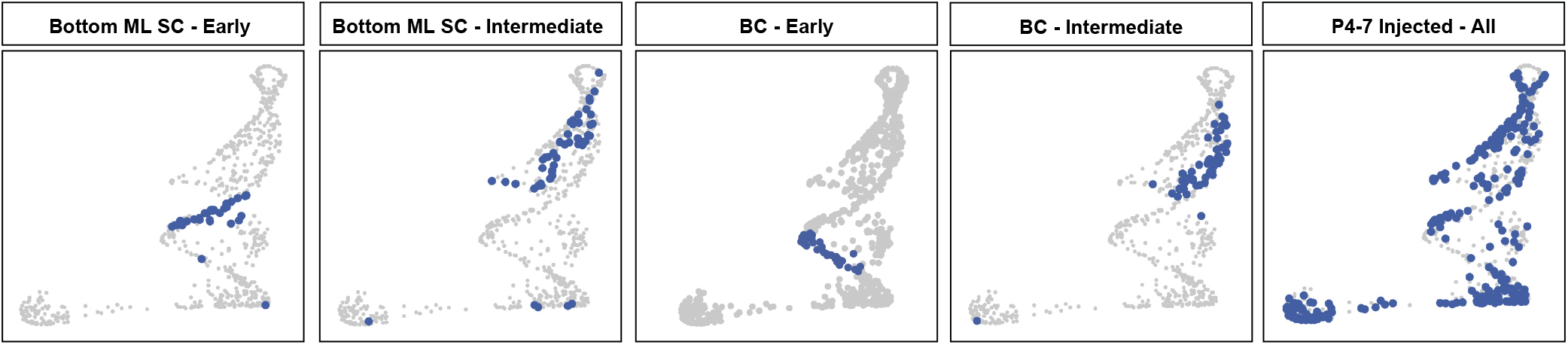
MLI line ages form discontinues s trajectories by Palantir. Early and intermediate bottom ML SCs represent cluster 5and 7 cells from Figure 6, respectively. Early and intermediate BCs depresent cluster 3and 6 cells from Figure 6, respectively. Early BCs, bottom MLSCs, and middle/uper ML SCs are largely non-ovealapping, suggesting subtype-specific clustering from the beginning of pseudotime.

**Supplementary Figure S10.**
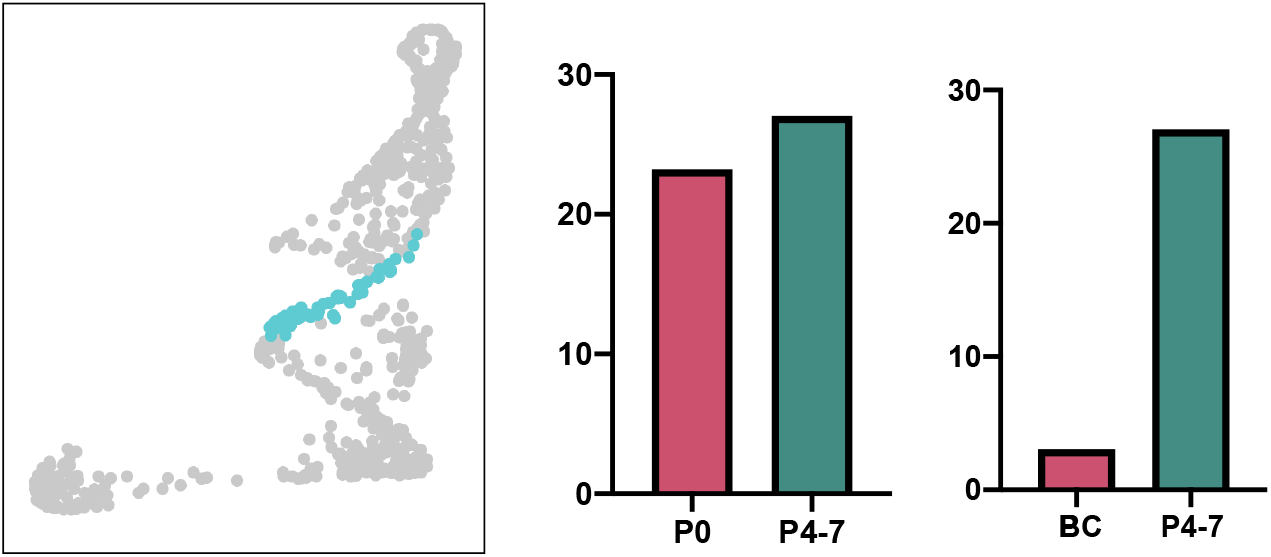
Cluster 6 contains early-born SC1s. Cluster 6 contained similar numbers of early- and late-born MLIs. However, this is due to the the high enrichment of SC1s within this cluster, as seen after removing SC1s from the early born population *(right).*

## Notes

### Competing Interest Statement

The authors have declared no competing interest.

### Summary of Updates

This version of the manuscript has been revised with updated discussion and supplemental files.

